# The acoustic properties, syllable structure, and syllable sequences of ultrasonic vocalizations (USVs) during neonatal opioid withdrawal in FVB/N mouse substrains

**DOI:** 10.1101/2024.11.06.622304

**Authors:** Kelly K. Wingfield, Teodora Misic, Sophia A. Miracle, Carly S. McDermott, Kaahini Jain, Nalia M. Abney, Kayla T. Richardson, Mia B. Rubman, Jacob A. Beierle, Elisha M. Wachman, Camron D. Bryant

## Abstract

Concomitant with the opioid epidemic, there has been a rise in pregnant women diagnosed with opioid use disorder and cases of infants born with neonatal opioid withdrawal syndrome (NOWS). NOWS refers to signs and symptoms following cessation of prenatal opioid exposure that comprise neurological, gastrointestinal, and autonomic system dysfunction. A critical indicator of NOWS severity is excessive, high-pitched crying. However, NOWS evaluation is, in large part, subjective, and additional cry features may not be easily recognized during clinical assessment. Thus, there is a need for more objective measures to determine NOWS severity. We used a third trimester-approximate opioid exposure paradigm to model NOWS traits in genetically similar inbred substrains of FVB/N mice (NJ, NCrl, NHsd, and NTac). Pups were injected twice daily from postnatal day 1 (P1) to P14 with morphine (10 mg/kg, s.c.) or saline (20 ml/g, s.c.). Because there were only very minor substrain differences in spontaneous withdrawal-induced ultrasonic vocalization (USV) profiles, we collapsed across substrains to evaluate the effects of morphine withdrawal on additional USV properties. We identified syllable sequences unique to morphine-withdrawn and saline-control FVB/N pups on P7 and P14. We also observed an effect of spontaneous morphine withdrawal on the acoustic properties of USVs and specific syllables on P7 and P14. Multiple withdrawal traits correlated with some acoustic properties of USVs and syllable type emission in morphine-withdrawn FVB/N pups on P7 and P14. These data provide an in-depth investigation of mouse USV syllable profiles and acoustic features during spontaneous neonatal opioid withdrawal in mice.

## 1. Introduction

Opioid use by pregnant individuals continues to be a public health concern, as misuse can lead to an increased risk of developing opioid dependence, overdose, and adverse pregnancy outcomes [1–3]. Specifically, opioid use during pregnancy exposes the fetus to opioids and can lead to infants developing neonatal opioid withdrawal syndrome (NOWS) after delivery [4–7]. NOWS is characterized by a set of withdrawal signs observed in infants at birth caused by the spontaneous cessation of prenatal opioid exposure and comprises gastrointestinal irregularities, autonomic nervous system dysfunction, and neurological dysregulation [8–13]. The onset, duration, and severity of NOWS features are highly variable, which can be attributed to various factors such as genetics, sex of the infant, gestational age of exposure, type and duration of opioid exposure, maternal polysubstance use, clinical care model, and environmental factors [14–26]. NOWS treatments include non-pharmacological interventions that promote maternal care, such as rooming-in and breastfeeding, and pharmacological treatment with mu-opioid receptor agonists (methadone, buprenorphine, morphine) [27,28].

Excessive crying is one of five critical indicators used to diagnose NOWS [29]. In human neonates, cries are scored using the Finnegan Neonatal Abstinence Scoring System (FNASS) or the Eat, Sleep, Care (ESC) assessment approach based on the level of support needed to console the infant [8,30,31]. NOWS infants who have excessive crying and who cannot be consoled despite comfort measures (swaddling, rocking, pacifier) are scored higher than infants who are easy to console [8,31,32]. Moreover, as part of the FNASS tool, clinicians must decide if cries are high-pitched, which is not explicitly defined [8,30]. Therefore, current methods of cry assessment are subjective and qualitative. Several studies have shown that infants prenatally exposed to opioids exhibit perturbations in cry frequencies and utterances that are not easily detectable by human listeners [33–36]. Thus, quantitative evaluation of the acoustic properties of cries may be useful for predicting NOWS severity and guiding treatment interventions.

Rodent models for NOWS traits are used to evaluate the effects of gestational opioid exposure on the neurobiological adaptations that contribute to withdrawal symptom onset, duration, and severity [37–39]. Rodent models permit control over multiple factors that contribute to variations in NOWS severity, such as environmental conditions, the type and duration of opioid exposure, maternal pregnancy co-exposures, and genetics. We use a third trimester approximate mouse model for NOWS that consists of morphine exposure from postnatal day (P) 1 to P14 [37–40]. This period in mice is functionally equivalent to the neurodevelopmental events that occur during the third trimester of human pregnancy [40,41], a period of exposure in humans necessary to observe NOWS [4]. Moreover, neonatal opioid exposure during the first two postnatal weeks is sufficient to produce withdrawal traits in mice [42–45]; thus, we can use a drug regimen comprising this exposure period to evaluate the neurobiological mechanisms underlying opioid withdrawal behaviors. Importantly, we can model withdrawal traits that are difficult to study in the clinic, such as irritability and excessive crying.

Neonatal mice emit ultrasonic vocalizations (USVs) when separated from their nest as a distress signal to promote maternal attention and rescue [46–52]. Furthermore, several studies have observed increased USVs during neonatal opioid withdrawal [42–44,53,54]. Thus, USVs can be utilized as a behavioral model for negative affective states, including opioid withdrawal-induced dysphoria. USVs are classified into syllables based on their spectrotemporal properties, frequency and duration [47,51,52,55,56]. In a previous study, we identified a unique USV signature associated with spontaneous morphine withdrawal on P14, defined by an increase in the percentage of Complex 3 syllables emitted, suggesting that specific components of USV profiles can reflect the severity of the aversive state of opioid withdrawal [45].

In this study, we used a third trimester-approximate mouse model for NOWS in four genetically similar inbred FVB/N substrains due to their reliable reproductive success and consistently large litter sizes [57]. Mouse substrains originate from the same parental strain (e.g. FVB/N, C57BL/6, BALB/c), but are separated from their parent colony for ≥ 20 generations, leading to the accumulation of spontaneous mutations and genetic drift [58]. Consequentially, this encroachment and fixation of alleles can result in phenotypic differences that can more readily be genetically mapped and validated on near-isogenic backgrounds [59–62].

We first sought to identify substrain differences in NOWS model traits, which would pave the way for future genetic mapping studies and identification of potential causal genes associated with withdrawal symptom severity. However, we did not observe robust substrain differences in opioid withdrawal traits. Thus, for a large portion of this study, we combined FVB/N substrains to increase our sample size and statistical power to deeply characterize the neonatal USV signatures reflecting spontaneous morphine withdrawal. We observed disruptions in the frequency, amplitude, and length of USVs akin to the cry characteristics observed in NOWS infants. Our detailed, objective analysis provides quantitative, descriptive results of complex acoustic features of withdrawal-induced USVs in a NOWS mouse model. Importantly, a similar approach could be implemented to delineate the cries of human NOWS infants and improve withdrawal symptom assessment.

## 2. Materials & Methods

### 2.1. Mice

All experiments involving mice were conducted following the National Institutes of Health *Guide for the Care and Use of Laboratory Animals* and were approved by the Institutional Animal Care and Use Committees at Boston University and Northeastern University. FVB/N inbred mice were purchased at 8 weeks old from various vendors and delivered together. FVB/NCrl: Charles River Laboratories (Strain #207); FVB/NHsd: Inotiv (Strain #118); FVB/NJ: The Jackson Laboratory (Strain #001800); FVB/NTac: Taconic Biosciences. All four substrains were included in each experimental cohort. Mice were provided *ad libitum* laboratory breeding chow (Teklad 18% Protein Diet, Envigo) and tap water and maintained on a 12 h light/dark cycle (lights on at 0630 h). Breeders of all four substrains were paired in-house after one week of acclimation to the vivarium. All breeder cages contained nestlets. Sires were removed seven days after breeder pairing to control for cage environment and avoid untimed pregnancies. Phenotyping occurred during spontaneous withdrawal (16 h) between 0900 h and 1100 h and was performed by female experimenters to control for the effect of experimenter sex on rodent behavior [63,64]. A portion of the FVB/NJ substrain data was previously published [45], thus substrain-collapsed analyses contain data from the FVB/NJ substrain. Substrain-interactive results reported in the Supplementary Material contain new traits not previously published (e.g., tail withdrawal latency and the acoustic properties of USVs and syllables), as well as published traits from the FVB/NJ substrain in the context of substrain differences (e.g., body weight, body temperature, hot plate latency and velocity, USV locomotor activity, USV emission, and syllable profiles).

### 2.2. Tattooing of mice

Pup tails were tattooed (ATS-3 General Rodent Tattoo System, AIMS) on postnatal day (P) 7 following behavioral testing and morning injections for identification and were returned to their home cage.

### 2.3. Morphine administration in FVB/NJ pups from P1 to P15

We used a third trimester-approximate mouse model of NOWS where genetically similar substrains of inbred FVB/N mice (NCrl, NHsd, NJ, NTac) pups were injected twice daily with either morphine (10 mg/kg) or saline (20 ml/g, s.c.) from postnatal day (P) one to P14 [42,44]. This mouse model is functionally equivalent to in-utero opioid exposure during the third trimester of human pregnancy [37–39,65]. Preclinical and clinical studies indicate that opioid exposure during the third trimester is both necessary and sufficient to induce a neonatal withdrawal state [40,41,66]. This model permits control of individual dosing and avoidance of maternal opioid exposure and potential negative consequences of maternal care and offspring behavior. Given that FVB/N substrains are nearly genetically identical (containing tens of thousands of variants compared to genetically diverse mice that contain millions of variants [67,68]) substrain differences in withdrawal-associated traits can allow for the identification of potential genomic loci containing causal gene variants associated with phenotypic differences [58,69]. Pups were sexed on P1, and each litter was approximately treatment- and sex-balanced to control for cage environment across treatments. From P1 – P15, injections of either morphine sulfate pentahydrate (10 mg/kg, 20 ml/kg, s.c.; Sigma-Aldrich) or saline (0.9%, 20 ml/kg, s.c.) were administered twice daily at 0900 h and 1700 h. Behavioral phenotyping occurred on P7 and P14 during spontaneous opioid withdrawal at 16 h post-morphine administration. On P7 and P14, morning injections were administered following phenotyping at approximately 1100 h.

### 2.4. Recording of ultrasonic vocalizations (USVs)

Pups were each placed into individual Plexiglass boxes (43 cm length x 20 cm width x 45 cm height; Lafayette Instruments) within a sound-attenuating chamber (Med Associates). USVs were recorded using the Ultrasound Recording Interface (Avisoft Bioacoustics UltrasoundGate 816H) for 10 min (P7) or 15 min (P14). Locomotor activity during all USV testing sessions was recorded using infrared cameras and tracked with ANY-maze software (Stoelting).

### 2.5. Thermal nociception testing on P7 and P14 in FVB/N pups

After USV recordings, pups were removed from the sound-attenuating chamber and placed in a Plexiglass cylinder (diameter, 15 cm; height, 33 cm) on a 52.5°C hot plate (IITC Life Science). On P7, the nociceptive response was defined as the latency for the pup to roll onto its back, an avoidance response that typically occurs as they attempt to lick their hind paw [42]. On P14, the nociceptive response was defined as the latency to jump, attempt to jump, hind paw lick or attempt to hind paw lick. Pups were removed from the hot plate immediately after observing a nociceptive response or after the 30 s cut-off (P7) or the 60 s cut-off (P14) if no pain response was observed.

Following hot plate testing, each pup was scruffed and the lower half of its tail was quickly lowered into a 48.5°C hot water bath (LX Immersion Circulator, PolyScience), and the latency (s) to withdraw the tail from the water was recorded. Pups were removed from the hot water bath immediately after observing the nociceptive response or after the 15 s cut-off if no response was observed. After nociceptive testing, each pup was weighed, their morning injection was administered, and they were returned to their home cage.

### 2.6. Supervised USV classification

DeepSqueak [70] and MATLAB (version 2022a) were used to detect individual USVs from mouse pup audio (.wav files) obtained from Avisoft microphones and software. USV call length (ms), principal frequency (kHz; median frequency), low frequency (kHz), high frequency (kHz), peak frequency (kHz; frequency with the greatest amplitude), frequency change (kHz; high – low frequency), and mean power (dB/Hz) were used to train a random forest classifier in Python. Additional information regarding syllable classification can be found in our previous study [45] and is available on our GitHub (https://github.com/camronbryant/NOWS_USV_classifier).

### 2.7. USV syllable repertoire randomness using Zipf’s law

Zipf’s law describes the complexity of language based on the frequency of words. It states that few words are used very frequently (e.g., “the”, “and”, “a”), while many words are rarely used [71,72]. Zipf’s law suggests an inverse relationship between words and their frequency, which reduces the effort required for communication by the speaker and listener (also known as the principle of least effort). We calculated the Zipf slope using the methods described in [55]. For each treatment and age, the logarithm of the number of each syllable was plotted against the logarithm of the syllable rank (descending order; the most common syllable rank = 1), where the slope of the line represents the Zipf value for each treatment/day syllable repertoire. A Zipf slope closer to 0 reflects diverse and random vocalization patterns, while a more negative Zipf slope depicts less diverse and repetitive vocalization patterns [55]

### 2.8. USV Sequences

Sequences were defined as USV bouts containing ≥ 3 syllables occurring with ≤ 30 ms separation, consistent with the literature regarding the temporal organization of mouse USVs [50,73]. We calculated the percentage of the top 10 most common sequences observed in morphine-withdrawn and saline-treated pups on P7 and P14 to compare treatment groups.

### 2.9. Exploratory factor analysis

We included 25 variables in factor analysis for P7 and P14, including body weight, body temperature, thermal algesia (hot plate latency, hot plate velocity, tail withdrawal latency), locomotor activity in isolation during USV recordings (average velocity and total distance), and 18 variables related to USV features and syllables. Exploratory factor analysis was performed using the R package “psych” and “fa” function. Variables were standardized to z scores before making the correlation matrices using the R package, “corrplot” with Pearson’s correlation coefficient. To determine the optimal number of factors to extract, we conducted a parallel analysis using the minimum residual (“minires”) method. Factors with eigenvalues greater than one were included in the analysis. The analysis revealed that five factors were sufficient. To simplify the factor loading matrix, we used the “varimax” function, which allows for easier interpretation of the relationships between variables.

### 2.10. Statistical analysis

Analysis was performed in R (https://www.r-project.org/). All data are presented as the mean ± standard error of the mean (SEM), and *p* < 0.05 was considered significant. Body weight, temperature, USV locomotion, and USVs over time were analyzed using linear mixed models (R package “lme4”) with Saline Treatment, Female Sex, and NCrl Substrain as the reference variables and Pup as a random effect for repeated measures. Sex and Substrain were removed from the model if there were no interactions. Significant interactions of interest (Morphine Treatment x Substrain, Morphine Treatment x Substrain x Sex) were followed up with least-square means (R package “emmeans”) using Tukey’s Honestly Significant Difference (HSD) tests. All other data were analyzed using linear models with Saline Treatment, Female Sex, and NCrl Substrain as the reference variables.

## 3. Results

### 3.1. Neonatal morphine exposure induces spontaneous withdrawal traits on P7 and P14 in four FVB/N substrains

The experimental timeline is provided in **Fig.1A**. Low birth weight and poor body temperature regulation are hallmark features of prenatal opioid exposure and NOWS in human infants [8,10]. We did not observe a Morphine Treatment x Substrain interaction on the following withdrawal traits, so the data were collapsed across substrain. Twice daily injections of morphine (10 mg/kg, s.c.) from P1 – P14 reduced body weight (**Fig.1B**) and temperature (**Fig.1C**) of pups from all four FVB/N substrains (NCrl, NHsd, NJ, NTac). Increased pain sensitivity (hyperalgesia) is frequently observed during morphine withdrawal in humans and mice [74,75]. During spontaneous withdrawal (16 h) on P7, morphine-withdrawn pups showed spontaneous thermal hyperalgesia, as indicated by a reduction in hot plate latency (**Fig.1D**). There was no difference in velocity during the time spent on the hot plate leading up to the nociceptive response, suggesting that locomotor activity did not confound the reduced hot plate latency in morphine-withdrawn pups (**Fig.1E**). Morphine-withdrawn pups also displayed decreased tail withdrawal latencies compared to saline control pups (**Fig.1F**), providing an additional measure of hyperalgesia.

**Figure 1.**
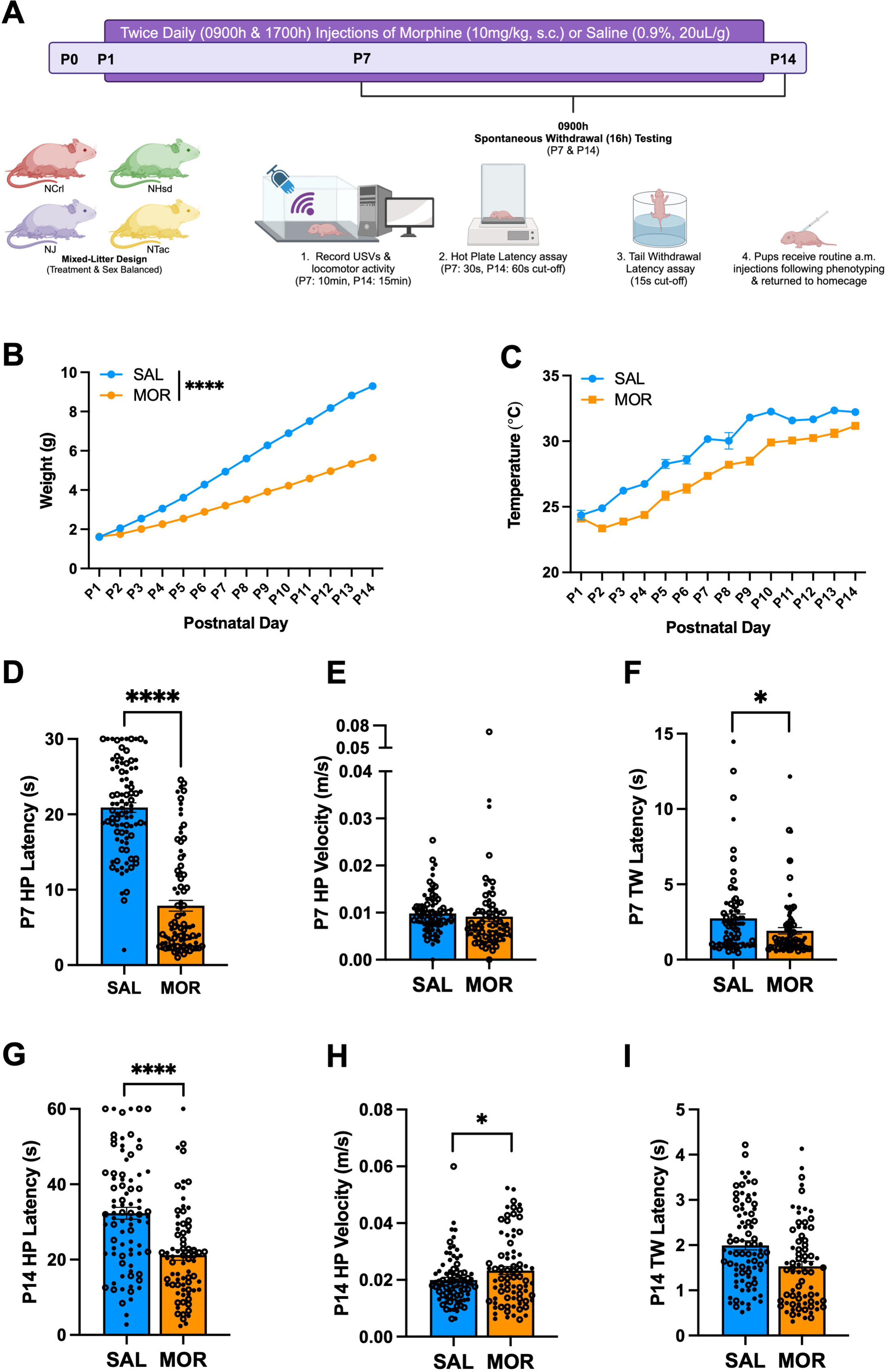
Morphine exposure from P1 – P14 is sufficient to induce opioid withdrawal traits in FVB/N pups. Data are plotted as the mean ± SEM. Closed circles = females. Open circles = males. Data were analyzed using linear mixed models with Saline Treatment, Female Sex, and NCrl Substrain as the reference variables and Pup as a random effect (for repeated measures). **A. Experimental Timeline. B. Body Weight:** The effect of Morphine Treatment on body weight was dependent on Postnatal Day (β = -0.29, SE = 0.012, t(166) = -24.14, \*\*\*\**p* < 0.0001), where morphine-withdrawn pups weighed significantly less than saline-control pups from P2 – P14 (P2: \*\**p* = 0.0027; P3 – P14: \*\*\*\**p* < 0.0001). **C. Temperature:** There were main effects of Morphine Treatment (β = -1.98, SE = 0.30, t(475) = -6.65, \*\*\*\**p* < 0.0001) and Postnatal Day (β = 0.65, SE = 0.041, t(2182) = 15.97, \*\*\*\**p* < 0.0001), but no interaction (β = 0.00042, SE = 0.026, t(2181) = 0.016, *p* = 0.99). SAL, n = 84; MOR, n = 84. **D. P7 Hot Plate Latency:** There were no interactions (all *p* ≥ 0.11). Morphine pups displayed decreased hot plate latency compared to saline-control pups (β = -13.03, SE = 0.95, t(177) = -13.70, \*\*\*\**p* < 0.0001). SAL, n = 90 (52F, 38M); MOR, n = 89 (43F, 46M). **E. P7 Hot Plate Velocity:** There were no interactions (all *p* ≥ 0.91). There was no effect of Morphine Treatment on hot plate velocity (β = -0.00072, SE = 0.0026, t(158) = -0.28, *p* = 0.78). SAL, n = 83 (48F, 35M) ; MOR, n = 79 (39F, 40M) . **F. P7 Tail Withdrawal Latency:** There were no interactions (all *p* ≥ 0.15). Morphine-withdrawn pups displayed reduced tail withdrawal latencies compared to saline-control pups (β = -0.76, SE = 0.34, t(173) = -2.25, \**p* = 0.026). SAL, n = 77 (44F, 33M); MOR, n = 86 (41F, 45M). **G. P14 Hot Plate Latency:** There were no interactions (all *p* ≥ 0.29). Morphine-withdrawn pups displayed reduced hot plate latencies compared to saline-control pups (β = -10.82, SE = 2.14, t(157) = -5.053, \*\*\*\**p* < 0.0001). SAL, n = 83 (48F, 35M; MOR, n = 75 (36F, 39M). **H. P14 Hot Plate Velocity:** There were no interactions (all *p* ≥ 0.63). Morphine-withdrawn pups displayed increased hot plate velocity compared to saline-control pups (β = 0.0033, SE = 0.0017, t(158) = 1.99, \**p* = 0.049). SAL, n = 83 (47F, 36M); MOR, n = 77 (37F, 40M). **I. P14 Tail Withdrawal Latency:** There were no interactions (all *p* ≥ 0.18). There was no effect of Morphine Treatment on tail withdrawal latency (β = -0.32, SE = 0.21, t(164) = -1.47, *p* = 0.14). SAL, n = 82 (48F, 34M); MOR, n = 75 (35F, 40M).

On P14, morphine-withdrawn pups once again displayed spontaneous thermal hyperalgesia on the hot plate (**Fig.1G**). Furthermore, morphine-withdrawn pups showed increased locomotor velocity on the hot plate on P14 (**Fig.1H**). Given that increased hot plate velocity would be expected to reduce contact time with the hot plate and thus, increase the latency to elicit a pain response, we can again conclude that locomotor activity did not confound the nociceptive response. There was no significant difference in tail withdrawal latency between morphine-withdrawn and saline control pups on P14 (**Fig.1I**).

### 3.2. Isolation-induced USV locomotor activity during spontaneous morphine withdrawal on P7 and P14 in FVB/N substrains

There were no Morphine x Substrain interactions on any of the following traits, so the data were collapsed across substrain. During spontaneous withdrawal on P7, all morphine-withdrawn FVB/N pups displayed increased locomotor activity compared to saline-treated pups in isolation during the USV recordings, specifically during the first 5 min (**Fig.2A**) and traveled a greater distance overall compared to saline controls (**Fig.2B**). There was no difference in the velocity while mobile between treatment groups (**Fig.2C**). There was no Morphine Treatment x Time interaction on USV distance traveled on P14 (**Fig.2D**). When summed over time, all morphine-withdrawn pups traveled a greater distance than saline pups (**Fig.2E**) and showed increased velocity (**Fig.2F**) on P14.

**Figure 2.**
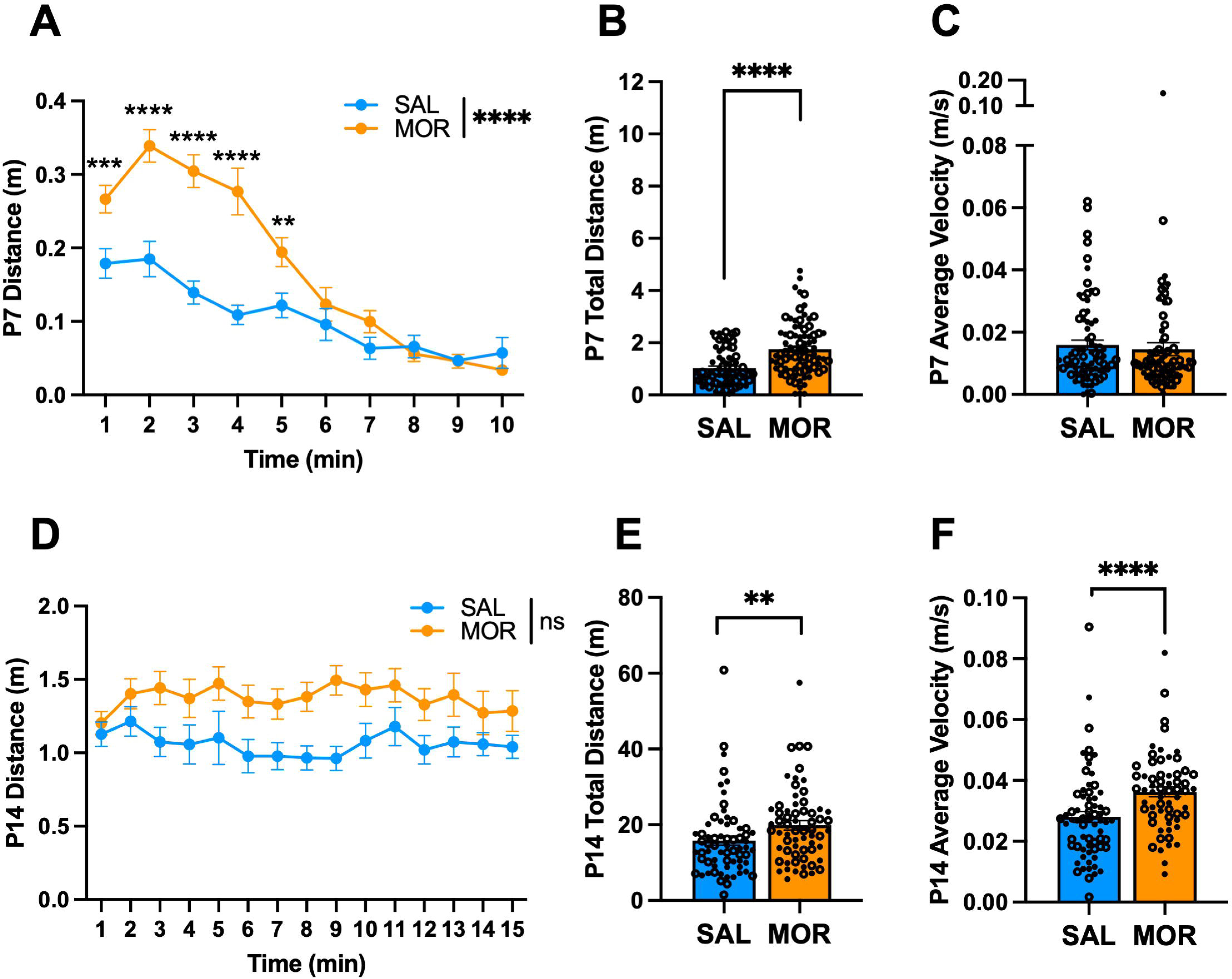
USV locomotor activity during spontaneous morphine withdrawal on P7 and P14 in FVB/N pups. Data are plotted as the mean ± SEM. Closed circles = females. Open circles = males. Data were analyzed using linear mixed models with Saline Treatment, Female Sex, and NCrl Substrain as the reference variables and Pup as a random effect (for repeated measures). **A. P7 USV Distance**: The effect of Morphine Treatment on USV distance was dependent on Time (β = -0.020, SE = 0.0026, t(1488) = -7.67, \*\*\*\**p* < 0.0001), where morphine-withdrawn pups traveled a greater distance than saline-control pups from 1 – 5 min of the recording session (all \*\**p* ≤ 0.0062). **B. P7 Total USV Distance**: There were no interactions (all *p* ≥ 0.57). Morphine Treatment was associated with a greater distance traveled during USV recordings associated with spontaneous opioid withdrawal (β = 0.68, SE = 0.14, t(165) = 4.98, \*\*\*\**p* < 0.0001). **C. P7 Average USV Velocity**: There were no interactions (all *p* ≥ 0.54). There was no effect of Morphine Treatment on average velocity (β = -0.00097, SE = 0.0025, t(167) = -0.39, *p* = 0.70). SAL, n = 78 (44F, 34M); MOR, n = 82 (43F, 39M). **D. P14 USV Distance**: There was no Morphine Treatment x Time interaction on USV distance (β = 0.005, SE = 0.014, t(1981) = 0.37, *p* = 0.71). **E. P14 Total USV Distance**: There were no interactions (all *p* ≥ 0.078). Overall, Morphine Treatment was associated with a greater distance traveled during USV recordings associated with spontaneous opioid withdrawal (β = 4.48, SE = 1.69, t(134) = 2.65, \*\**p* = 0.0091). **F. P14 Average USV Velocity**: There were no interactions (all *p* ≥ 0.63). Morphine Treatment was associated with increased average velocity (β = 0.0083, SE = 0.0025, t(133) = 3.39, \*\*\**p* = 0.00092). SAL, n = 67 (37F, 30M) MOR, n = 62 (29F, 33M)

### 3.3. USV emission profiles during spontaneous morphine withdrawal on P7 in FVB/N substrains

USVs are emitted by neonatal mice in isolation to communicate a negative internal state and promote maternal attention [48,49,76]. Thus, USVs can model irritability and excessive crying observed in infants. There was no Morphine Treatment x Substrain x Time interaction on the following traits, so we collapsed across substrains. During spontaneous withdrawal (16 h) on P7, morphine-withdrawn FVB/N pups emitted more USVs during the first 4 min of the recording session compared to saline pups. In contrast, during the 6 – 10 min time interval, morphine-withdrawn pups emitted fewer USVs compared to saline pups (**Fig.3A**). When summed over time, there was no difference in total USVs collapsed across the four FVB/N substrains (**Fig.3B**). However, there was a Morphine Treatment x Sex x Substrain interaction driven by the NTac substrain, where morphine-withdrawn females emitted more USVs than saline control females (**Fig. S1A**).

**Figure 3.**
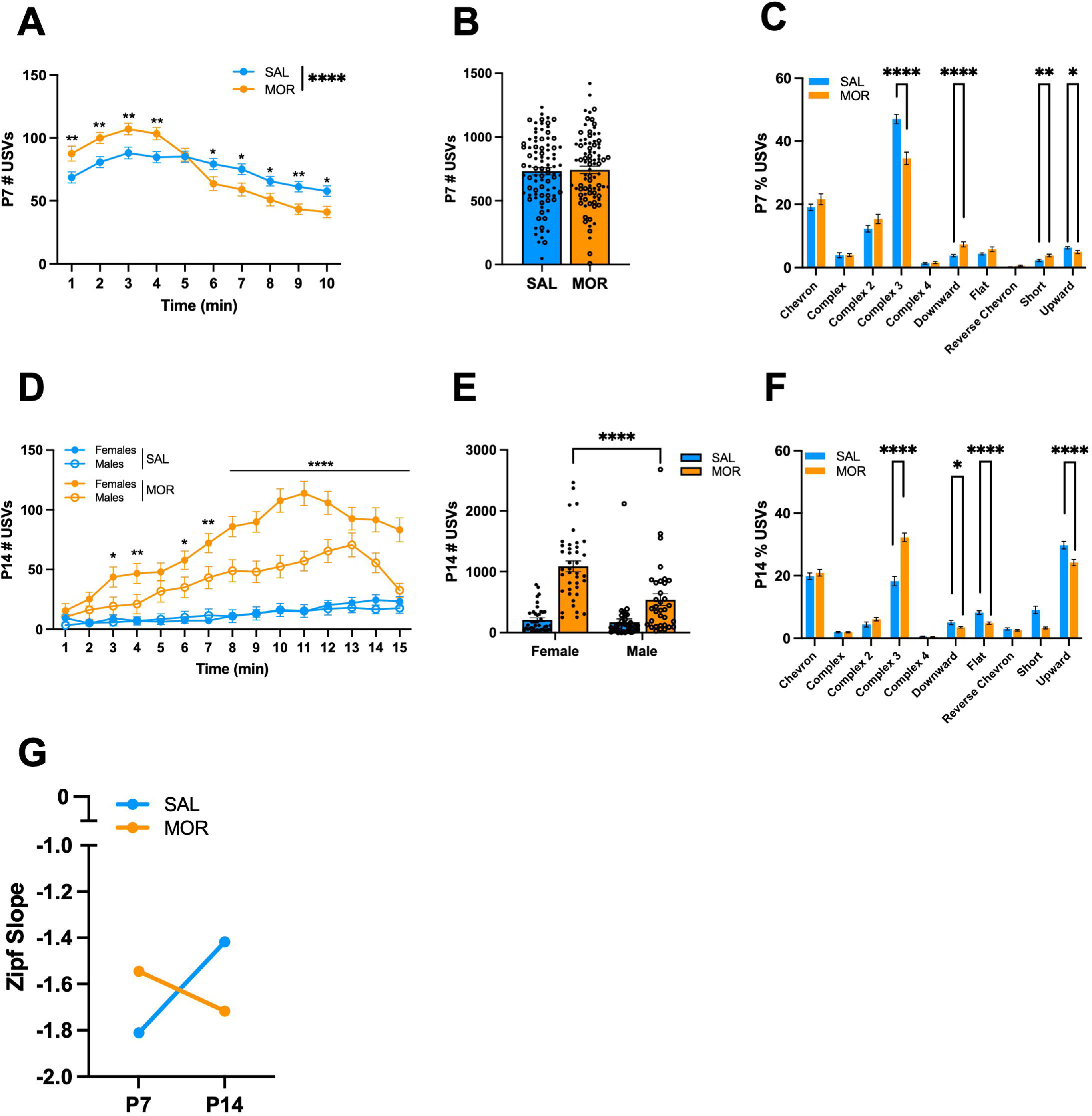
USV emissions during spontaneous morphine withdrawal on P7 in FVB/N pups. Data are plotted as the mean ± SEM. Closed circles = females. Open circles = males. Data were analyzed using linear mixed models with Saline Treatment, Female Sex, and NCrl Substrain as the reference variables and Pup as a random effect (for repeated measures). The following data are collapsed across Substrain and Sex for simplicity. **A. P7 USVs across time**: The effect of Morphine Treatment was dependent on Time (β = -5.38, SE = 0.62, t(1573) = -8.63, \*\*\*\**p* < 0.001, where Morphine Treatment was associated with an increase in USVs during the first 4 min of the recording session (all \*\**p* < 0.0055), and a decrease in USVs during the 6 – 10 min time intervals (all \**p* < 0.040). **B. P7 Total USVs**: Overall, there was no effect of Morphine Treatment on the total number of USVs (β = -0.99, SE = 40.64, t(181) = -0.024, *p* = 0.98). **C. P7 Syllable Profile**: Morphine Treatment was associated with a decrease in Complex 3 (β = -0.12, SE = 0.025, t(181) = - 5.04, \*\*\*\**p* < 0.001) and Upward (β = - 0.014, SE = 0.006, t(181) = -2.28, \**p* = 0.024), and an increase in Downward (β = 0.036, SE = 0.0089, t(181) = 4.04, \*\*\*\**p* < 0.0001) and Short (β = 0.015, SE = 0.0057, t(181) = 2.71, \*\**p* = 0.0074). **D. P14 USVs across time**: There was a Morphine Treatment x Sex x Time interaction (β = -2.32, SE = 0.53, t(2170) = -4.37, \*\*\*\**p* < 0.0001). Both morphine-withdrawn females and males vocalized more over time compared to saline-control pups; however, morphine-withdrawn females vocalized more than morphine-withdrawn males during the 3, 4, and 6 – 15 min intervals (all \**p* ≤ 0.015). **E. P14 Total USVs**: There was a Morphine Treatment x Sex interaction (β = -452.5, SE = 153.5, t(151) = -2.95, \*\**p* = 0.0037). Both morphine-withdrawn females (β = 883.63, SE = 93.49, t(80) 9.45, \*\*\*\**p* < 0.0001) and morphine-withdrawn males (β = 431.1, SE = 124.2, t(71) = 3.47, \*\*\**p* = 0.00089) vocalized more than saline-control pups. Furthermore, morphine-withdrawn females emitted more USVs than morphine-withdrawn males (β = 471.6, SE = 109, t(151) = 4.32, \*\*\*\**p* < 0.0001). **F. P14 Syllable Profile**: Morphine Treatment was associated with an increase in the proportion of Complex 3 syllables emitted during spontaneous opioid withdrawal (β = 0.14, SE = 0.20, t(155) = 6.89, \*\*\*\**p* < 0.0001), and a decrease in Downward (β = -0.015, SE = 0.0077, t(155) = -2.02, \**p* = 0.045), Flat (β = -0.034, SE = 0.0071, t(155) = -4.73, \*\*\*\**p* < 0.0001), Short (β = -0.058, SE = 0.013, t(155) = -4.61, \*\*\*\**p* < 0.0001), and Upward (β = -0.056, SE = 0.016, t(155) = -3.48, \*\*\**p* = 0.00065) syllables. **G. Zipf Slope**: The Zipf slope of morphine-withdrawn pups decreased from P7 to P14, demonstrating that the syllable repertoire became less random and more repetitious. The Zipf slope of saline-treated pups increased from P7 to P14, depicting increased syllable usage diversity. P7: SAL, n = 88 (50F, 38M); MOR, n = 87 (45F, 42M). P14: SAL, n = 79 (44F, 25M); MOR, n = 76 (38F, 38M).

Neonatal mouse USVs are classified into distinct syllables based on their spectrotemporal properties [51,52,55,56]. In a recent study, we identified a unique USV syllable profile associated with spontaneous morphine withdrawal in FVB/NJ mouse pups [45]. When collapsed across substrains, morphine-withdrawn pups emitted fewer Complex 3 and Upward syllables and a greater percentage of Downward and Short syllables than saline-treated pups on P7 (**Fig.3C**). We also observed interactions of Morphine Treatment with Sex and Substrain for Chevron, Complex 4, and Reverse Chevron syllables emitted (**Fig.S1B – D**).

### 3.4. USV emission profiles during spontaneous morphine withdrawal on P14 in FVB/N substrains

During spontaneous withdrawal on P14 (16 h), all morphine-withdrawn FVB/N pups of both sexes vocalized more than saline-treated pups during the 15 min USV recording session; however, the effect of Morphine Treatment was stronger in females, where morphine-withdrawn females emitted significantly more USVs than morphine-withdrawn males (**Fig.3D**). When summed over time and collapsed across substrains, morphine-withdrawn pups of both sexes showed a robust increase in USVs compared to saline pups. Again, morphine-withdrawn females vocalized approximately twice as much as morphine-withdrawn males (**Fig.3E**). Opposite to our P7 data, for P14, we observed a robust increase in the percentage of Complex 3 syllables emitted by all FVB/N morphine-withdrawn pups, as well as a decrease in Downward, Flat, Short, and Upward syllables (**Fig.3F**). Additionally, we observed Morphine Treatment x Sex x Substrain interactions on the proportion of Downward (**Fig.S2A**) and Upward (**Fig.S2B**) syllables emitted during spontaneous morphine withdrawal. We also calculated the Zipf slope in morphine-withdrawn and saline-treated pups on P7 and P14 to determine the diversity of syllable repertoire. On P7, the Zipf slope was closer to zero in morphine-withdrawn pups compared to saline-treated pups, indicating that morphine-withdrawn pups produce a more diverse syllable repertoire than saline-treated pups (**Fig.3G**). On P14, the Zipf slope was closer to –2, indicating a switch in that morphine-withdrawn pups emitted a more repetitive and less random syllable profile (**Fig.3G**).

### 3.5. Morphine withdrawal alters USV syllable sequences on P7 and P14 in FVB/N pups

Next, we explored USV syllable sequences to determine if morphine withdrawal was associated with a unique temporal syllable pattern On P7, we did not observe a Morphine Treatment x Substrain interaction in the total number of syllable sequences; therefore, we combined substrain USV data across substrains to provide additional power to determine whether a syllable sequence was associated with spontaneous withdrawal. On P7, there was no effect of Morphine Treatment on the total number of unique sequences emitted (**Fig.4A**). On P14, morphine-withdrawn pups emitted more unique syllable sequences than saline-treated pups (**Fig.4B**). Additionally, morphine-withdrawn females emitted significantly more unique sequences than morphine-withdrawn males. However, most sequences were emitted only once or infrequently on P7 and P14. Thus, we calculated the percentage of sequences within the top 10 most common sequences observed in morphine-withdrawn and saline-treated pups. On P7 and P14, we observed different syllable sequences unique to both treatment groups. On P7, morphine-withdrawn pups emitted a large percentage of sequences containing 3 – 5 Chevron (Ch) syllables, while saline-treated pups emitted a large percentage of sequences containing 3 – 5 Complex 3 (C3) syllables (**Fig.4C**). On P14, morphine-withdrawn pups emitted sequences containing four C3 syllables, C3 – C3 – Upward (Up) and C3 – Ch – Ch, which were absent in saline-treated pups (**Fig.4D**). The sequences, Ch – Up – Up and Up – C3 – Up, were only observed in saline-treated pups.

**Figure 4.**
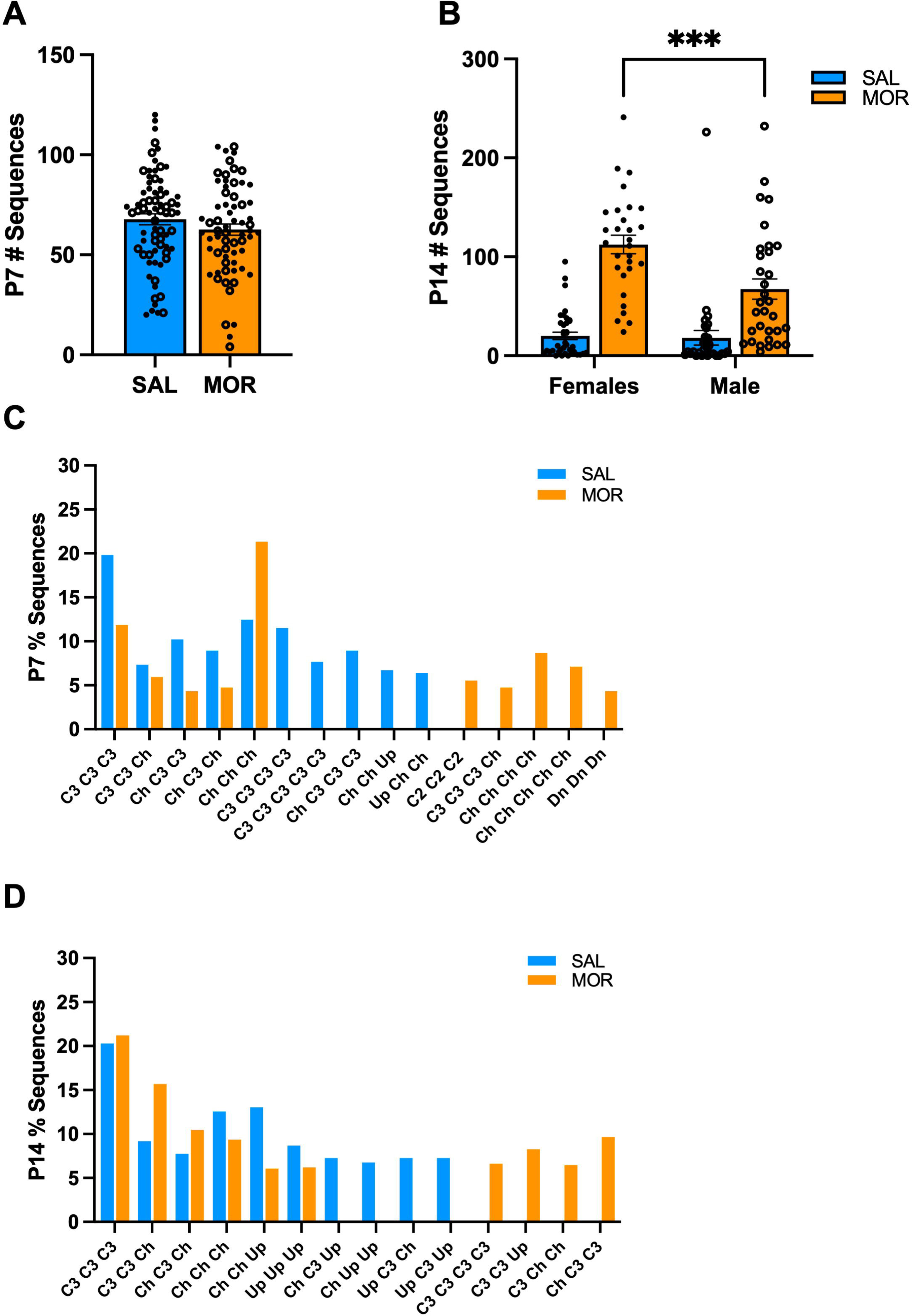
Unique USV syllable sequences in FVB/N pups on P7 and P14. Data are plotted as the mean ± SEM. Closed circles = females. Open circles = males. Data were collapsed across Substrain to increase statistical power and were analyzed using linear mixed models with Saline Treatment and Female Sex as the reference variables. **A. P7 Total USV Sequences**: There was no effect of Morphine Treatment on the total number of sequences emitted (β = -5.15, SE = 3.96, t(136) = -1.30, *p* = 0.20). SAL = 74 (40F, 34M); MOR = 65 (37F, 28M). **B. P14 Total USV Sequences**: There was a Morphine Treatment x Sex Interaction (β = 43.22, SE = 15.60, t(126) = 2.77, \*\**p* = 0.00065) where morphine-withdrawn females emitted more unique sequences than males (β = 45.20, SE = 11.31, t(126) = 3.89, \*\*\**p* = 0.0001). The following data is presented as the percentage of sequences of the top 10 most common sequences observed in morphine-withdrawn and saline-treated pups on **C. P7 % Sequences.** and **D. P14 % Sequences**. Complex 2 (C2), Complex 3 (C3), Chevron (Ch), Down (Dn), Upward (Up). SAL, n = 69 (38F, 31M); MOR, n = 61 (29F, 32M).

### 3.6. Morphine withdrawal alters the acoustic features of USVs during spontaneous withdrawal on P7

In humans, pitch (frequency), loudness (amplitude), and word length are vocal features necessary for emotional communication [77]. Moreover, several studies have observed altered affective communication in individuals with autism and schizophrenia, in addition to altered USV features in rodent models for these disorders [52,78–81]. Thus, we evaluated the spectrotemporal properties of vocalizations on P7 to further identify features that could reflect the negative affective state associated with spontaneous morphine withdrawal. When collapsed across substrains, morphine-withdrawn pups emitted USVs with reduced length (**Fig.5A**) and power compared to saline-treated pups (**Fig.5B**). Morphine-withdrawn pups emitted USVs with a higher low frequency, smaller change in frequency, and increased principal and peak frequencies compared to saline control pups (**Fig.5C**). There was also a Morphine Treatment x Sex x Substrain interaction on USV power (**Fig.S1E**).

**Figure 5.**
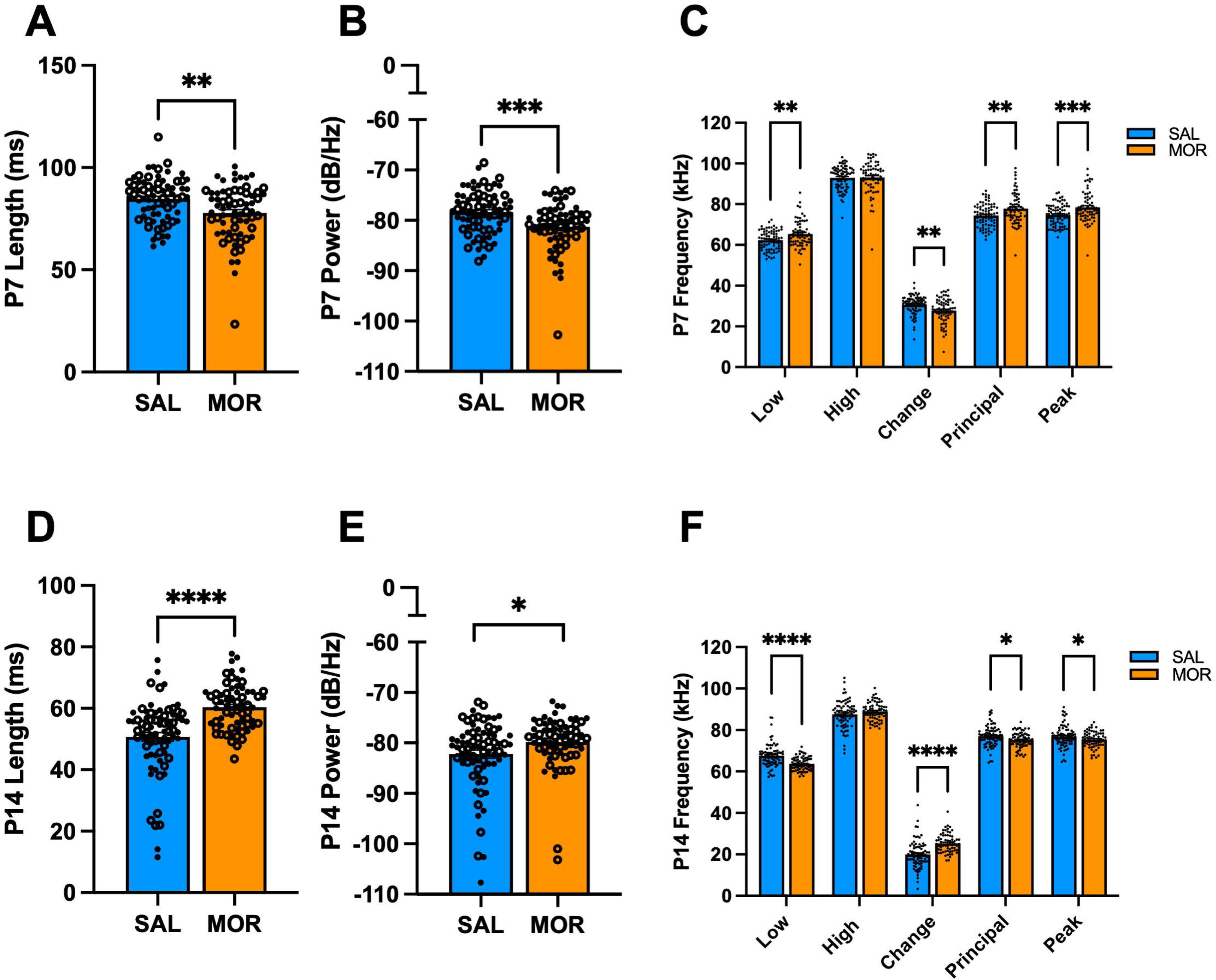
Acoustic features of USVs emitted during spontaneous morphine withdrawal in FVB/N pups on P7 and P14. Data are plotted as the mean ± SEM. For A,B, D, and E: closed circles = females; open circles = males. For C and F, closed circles include both sexes to avoid clutter. Data were analyzed using linear mixed models with Saline Treatment, Female Sex, and NCrl Substrain as the reference variables. The following data are collapsed across Substrain for simplicity. **A. P7 Length:** There were no interactions (all *p* ≥ 0.63). Morphine Treatment was associated with shorter USV duration (β = -0.0063, SE = 0.0021, t(144) = -3.054, \*\**p* = 0.0027). **B. P7 Power**: Morphine Treatment was associated with reduced USV power (β = -2.69, SE = 0.76, t(144) = -3.54, \*\*\**p* = 0.00055). **C. P7 Frequencies**: Morphine Treatment was associated with higher lowest frequency (β = 3.056, SE = 0.92, t(144) = 3.31, \*\**p* = 0.0012), reduced change in frequency (β = -2.94, SE = 0.88, t(144) = -3.34, \*\**p* = 0.0011), increased principal frequency (β = 3.36, SE = 1.08, t(144) = 3.12, \*\**p* = 0.0022), and increased peak frequency (β = 3.71, SE = 1.03, t(144) = 3.61, \*\*\**p* = 0.00042). There was no effect of Morphine Treatment on high frequency (β = 0.12, SE = 1.13, t(44) = 0.11, *p* = 0.92). SAL, n = 73 (41F, 32M); MOR, n = 66 (37F, 29M). **D. P14 Length**: There were no interactions (all *p* ≥ 0.091). Morphine Treatment was associated with increased USV duration (β = 0.010, SE = 0.0017, t(145) = 6.18, \*\*\*\**p* < 0.0001). **E. P14 Power**: There were no interactions (all *p* ≥ 0.36). Morphine Treatment was associated with increased power (β = 2.32, SE = 0.97, t(145) = 2.39, \**p* = 0.018). **F. P14 Frequencies**: Morphine Treatment was associated with reduced low frequency (β = -3.38, SE = 0.79, t(145) = -4.27, \*\*\*\**p* < 0.0001), a greater change in frequency (β = 5.41, SE = 0.93, t(145) = 4.04, \*\*\*\**p* < 0.0001), reduced principal frequency (β = -.89, SE = 0.73, t(145) = -2.59, \**p* = 0.011), and reduced peak frequency (β = -1.72, SE = 0.77, t(145) = -2.23, \**p* = 0.027). There was no effect of Morphine Treatment on high frequency (β = 1.43, SE = 0.93, t(145) = 1.54, *p* = 0.13). SAL, n = 75 (42F, 33M); MOR, n = 67 (33F, 34M).

Next, we assessed the acoustic features of USV syllables during spontaneous withdrawal on P7 to determine if specific syllable types drove the morphine withdrawal-induced alterations in USV features. There were Morphine Treatment interactions with Sex and/or Substrain on the acoustic features of Downward (**Fig.S3A–E**), Complex 4 (**Fig.S3F**), Flat (**Fig.S3G**), and Upward (**Fig.S3I**) syllables. When collapsed across Substrain, we observed alterations in length, power, and frequency of nearly all syllable types on P7 (**Fig.S4**).

### 3.7. Morphine withdrawal alters the acoustic features of USVs during spontaneous withdrawal on P14

On P14, morphine-withdrawn pups (substrain-collapsed) emitted longer USVs (**Fig.5D**) and increased power (**Fig.5E**) during spontaneous withdrawal compared to saline control pups. Morphine-withdrawn pups emitted USVs with reduced low frequency, increased frequency change, and reduced principal and peak frequency (**Fig. 5F**). There were Morphine Treatment interactions with Sex and/or Substrain on the acoustic features of Chevron (**Fig.S5A–C**), Complex (**Fig.S5D**), Complex 4 (**Fig.S5E**), Reverse Chevron (**Fig.S5F**), and Upward (**Fig.S5G**) syllables. We also observed Morphine Treatment x Sex x Substrain interactions on the low (**Fig.S6A**), high (**Fig.S6B**), and principal (**Fig.S6C**) frequencies of Complex 3 syllables during spontaneous withdrawal on P14. When collapsed across substrain, we observed fewer effects of morphine withdrawal on changes in syllable properties compared to saline controls. Only a few acoustic features of Reverse Chevron, Complex 3, and Downward syllables were affected during spontaneous morphine withdrawal (**Fig.S7**).

### 3.8. Exploratory factor analysis of USV features with withdrawal traits on P7 and P14

Spectrotemporal features of USVs (changes in frequency, length, and power) may be useful for communicating different aversive states in mice [82–84]. Thus, we aimed to determine if certain acoustic properties of USVs or syllable emissions correlated with the severity of other withdrawal traits. On P7, for saline-treated pups, USV features and syllables did not load onto any shared factors with other withdrawal traits (body weight and temperature, hyperalgesia, increased locomotor activity) in saline-treated pups (**Table S1**). However, for P7 morphine-withdrawn pups, tail withdrawal latency and % Complex syllables loaded onto a shared factor (**Table S2**). On P14, for saline-treated pups, hot plate velocity, USV length, USV power, % Chevron and % Short emissions loaded onto a common factor (**Table S3**). For P14 morphine-withdrawn pups, hot plate latency, tail withdrawal latency, and % Complex 2 emission loaded on the same factor (**Table S4**).

### 3.9. Correlational analysis of USV features with withdrawal traits on P7 and P14

On P7 (**Fig.6A**), in morphine-withdrawn pups (collapsed across substrain), lower body weight (a suspected indicator of a more severe response to repeated morphine treatment) correlated with reduced USVs, shorter USV length, reduced percentage of Chevron syllables, and increased percentage of Reverse Chevron syllables. Reduced hot plate latency (hyperalgesia) in morphine-withdrawn pups correlated with reduced percentage of Chevron syllables, while reduced tail withdrawal latency (hyperalgesia) correlated with reduced percentage of Complex, Complex 3, and Reverse Chevron syllables. Additionally, increased total distance traveled during USV recording correlated with a greater percentage of Upward syllables emitted.

**Figure 6.**
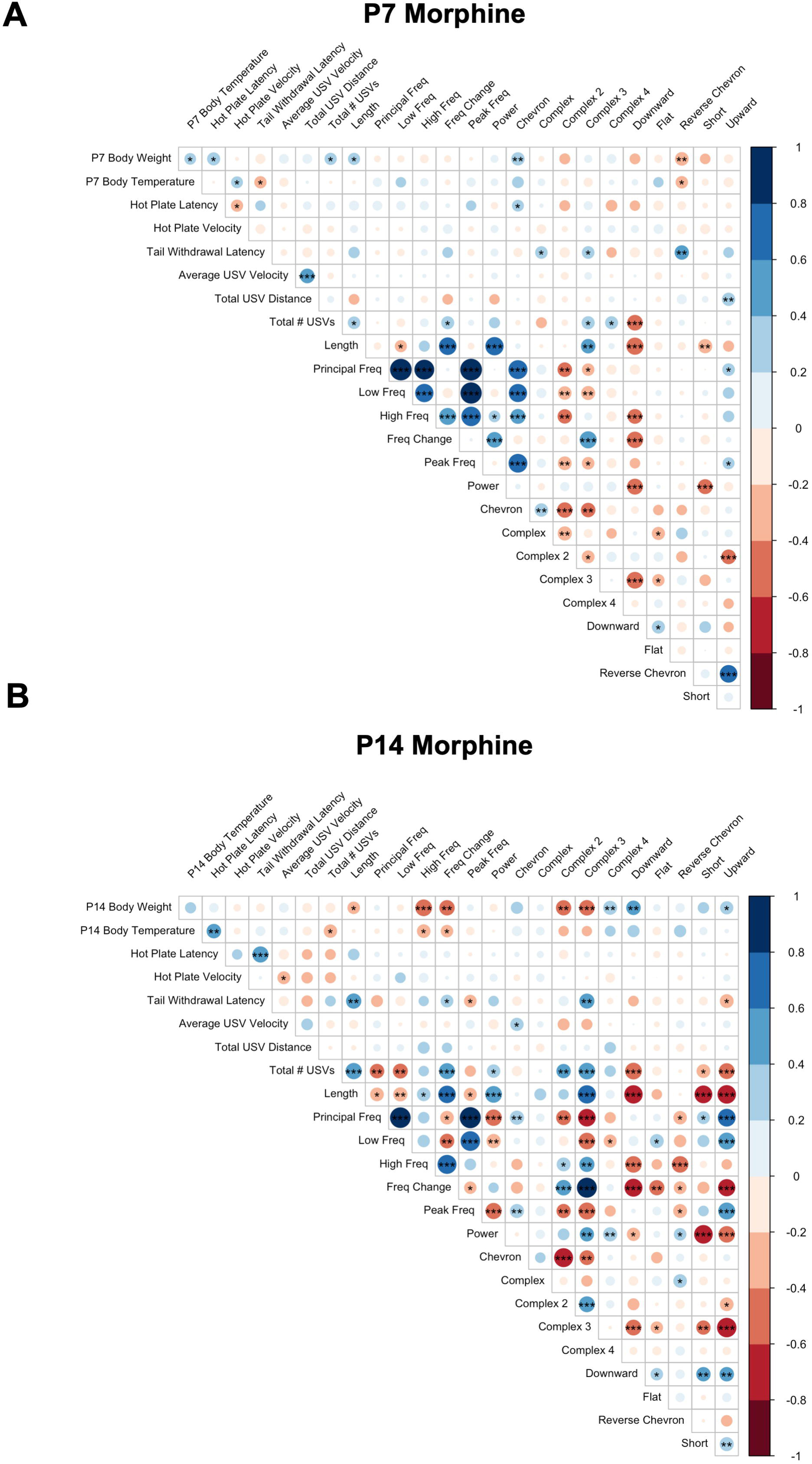
Correlation of withdrawal traits in morphine-withdrawn FVB/N pups on P7 and P14. Colors indicate Pearson’s correlation coefficient: blue = positive, red = negative. Darker colors reflect stronger correlations. \**p* < 0.01, \*\**p* < 0.001, \*\*\**p* < 0.0001. P7, n = 54 (28F, 26M); P14, n = 44 (21F, 23M).

On P14 (**Fig.6B**) in morphine-withdrawn pups, lower body weight correlated with longer USV length, greater high frequency, greater frequency change, increased percentage of Complex 2 and Complex 3 syllable emissions, and reduced percentage of Complex 4, Downward, and Upward syllable emissions. Reduced body temperature (a more pronounced physiological response to repeated morphine) correlated with increased USVs and greater frequency and frequency change. Reduced tail withdrawal latency (hyperalgesia) correlated with shorter USV length, reduced frequency change, greater peak frequency, reduced percentage of Complex 3 syllables, and increased percentage of Upward syllables.

In saline-treated pups, on P7 (**Fig. S8A**), low body weight also correlated with reduced USVs and increased percentage of Reverse Chevron syllables. Reduced hot plate latency (hyperalgesia) correlated with increased USV power, which was absent in morphine-withdrawn pups on P7. In contrast to morphine-withdrawn pups, reduced tail withdrawal latency in saline-treated pups correlated with increased percentage of Complex 3 syllables. On P14 (**Fig. S8B**), increased hot plate velocity correlated with reduced length, reduced low and high frequencies, reduced power, and reduced percentage of Complex 3 syllables in saline-treated pups, which was not observed in morphine-withdrawn pups on P14.

## 4. Discussion

A regimen of twice-daily injections of morphine from P1 – P14 was sufficient to induce opioid withdrawal traits in genetically similar inbred FVB/N mouse substrains (NCrl, NHsd, NJ, NTac), such as low body weight, hypothermia, increased pain sensitivity, enhanced USV emission, and altered USV syllable profiles. We first sought to identify substrain differences in NOWS model traits during morphine withdrawal with the long-term goal of conducting a future genetic mapping study in a reduced complexity cross [58,69]. However, we did not find robust trait differences across the four substrains. All FVB/N pups displayed similar withdrawal-induced USV profiles, including a robust increase in the percentage of Complex 3 syllables on P14, which we hypothesize is a biobehavioral marker associated with the severe negative internal state of spontaneous morphine withdrawal [45]. The similar results across substrains were a fortuitous opportunity to collapse the data across substrains to improve our power to detect the effects of morphine withdrawal on other USV features, such as syllable repertoire, syllable sequences and acoustic properties.

We calculated the Zipf slope for syllable profiles to determine if the withdrawal-induced USV profile contained significant information compared to the saline control profile. Interestingly, in morphine-withdrawn pups, the Zipf slope of the syllable repertoire was more negative than that of saline-treated pups on P14. A more negative Zipf slope indicates a less random repertoire, suggesting that pups can communicate meaningful information regarding withdrawal-induced dysphoria through repetitive syllables. In saline-treated pups, the Zipf slope decreased from P7 to P14, consistent with normal syllable repertoire development in mice [55]. Given the increase in the Zipf slope from P7 to P14 in morphine-withdrawn mice, we hypothesize that this change may be attributed to repetitive Complex 3 syllables during spontaneous morphine withdrawal.

High-pitched, excessive crying is a defining withdrawal symptom observed in human infants exposed to opioids and is one of the most important indicators used to assess NOWS severity in the hospital [29,33,34,36,85]. However, the FNASS or ESC evaluation approaches do not define a high-pitched cry, introducing the possibility of inconsistent evaluations across observers [8]. In a recent study by Manigualt et al., disruptions in additional acoustic properties of cries were observed in infants with NOWS compared to infants without NOWS [35]. These disruptions include changes in the fundamental frequency, frequency formants (frequency peaks due to vocal resonance), cry utterances (duration during respiratory expiration), and amplitude (power), which are undetectable by humans. Moreover, alterations in USV emission rate, syllable profiles, and frequency properties have been observed in rodent models for autism and schizophrenia [52,80,81]. These studies suggest that an unbiased, quantitative measurement of acoustic properties may provide an objective evaluation of affective dysregulation and withdrawal symptom severity beyond current assessment methods.

We observed higher principal (median) USV frequencies in morphine-withdrawn pups than saline controls on P7. In contrast, on P14, morphine-withdrawn pups emitted USVs with lower principal frequencies compared to saline controls. On P14 but not P7, morphine-withdrawn pups emitted USVs with increased power compared to saline control pups. Additionally, the peak frequency (the frequency with the highest power) was lower in morphine-withdrawn pups compared to saline pups. Thus, morphine-withdrawn pups emphasize lower (and louder) USV frequencies compared to saline pups. Moreover, Complex 3 syllables have the lowest principal frequency compared to other syllable types, further supporting its connection to a negative internal state. Interestingly, Lefebvre et al. observed an increased rate of low frequency USVs (≤ 60 kHz) emitted during restraint stress in adult mice [86]. Changes in acoustic properties have been observed primarily in lower frequency USVs in neonatal mice in response to environmental conditions such as low temperatures, isolation, and male odor [46]. Additionally, Ehret & Haack observed that USVs occurring within the low 20 – 60 kHz range promoted mouse pup retrieval in dams [87,88]. Together, these observations suggest that lower USV frequencies may indicate severe distress associated with spontaneous morphine withdrawal, thereby increasing maternal responsiveness.

Increased duration of the first cry utterance (initial vocalization) has also been observed in infants prenatally exposed to opioids [33,34]. On P7, there was no significant difference in USV length between treatment groups. On P14, morphine-withdrawn pups emitted longer USVs than saline control pups. However, when investigating the properties of individual syllables, only the length of Reverse Chevron syllables was greater in morphine-withdrawn pups compared to saline control pups, suggesting that the withdrawal-induced increase in overall USV length could be explained by an increased percentage of longer syllables (e.g., Complex 2, 3, and 4) emitted during spontaneous morphine withdrawal. We also considered the length of the initial syllable emitted during a sequence as a model for the first cry utterance. On P14, the most common, unique syllable sequences began with a Complex 3 syllable in morphine-withdrawn pups, which is longer in duration than Upward syllables, the initial syllable in common saline control sequences.

We evaluated USV syllable sequences to determine if certain orders of syllables were associated with morphine withdrawal. Overall, there was no significant treatment difference in the number of unique syllable sequences emitted on P7. In contrast, morphine-withdrawn pups emitted more unique sequences than saline control on P14. However, despite the numerous unique syllable sequences, most sequences occurred only once in both treatment groups. When investigating the percentage of the top 10 most common sequences, we identified a few unique sequences to morphine-withdrawn and saline-treated pups on P7 and P14. This could suggest that certain syllable sequences are emitted during different levels of distress (i.e. isolation-induced distress on top of spontaneous morphine withdrawal vs. isolation-induced distress alone in saline control pups). Conversely, these treatment differences could partly be explained by individual variation in syllable organization and potential random or non-meaningful sequences [55].

We observed correlations between USV features and other withdrawal traits on P7 and P14. On P7, lower body weight correlated with decreased USVs and shorter USVs in morphine-withdrawn pups. The positive correlation between body weight and USV emissions was also observed in saline-treated pups on P7, suggesting that this association may be due to immature respiratory structures unrelated to neonatal morphine exposure during development. Interestingly, reduced tail withdrawal latency (hyperalgesia) on P7 was associated with decreased Complex 3 emissions in morphine-withdrawn pups but increased Complex 3 emissions in saline-treated pups. However, on P14, when we normally observe more robust withdrawal phenotypes, reduced tail withdrawal latency continued to be associated with decreased Complex 3 emission in morphine-withdrawn pups, while there was no significant correlation in saline-treated pups. Other nociception phenotypes (hot plate latency and velocity) did not correlate with increased Complex 3 emissions in morphine-withdrawn and saline-treated pups on P14, suggesting that Complex 3 emission does not reflect sensitivity to stimulus-evoked pain. Additionally, lower body weight (a commonly observed physiological adaptation to chronic opioid administration in adult rodents) correlated with increased Complex 3 emissions only in morphine-withdrawn pups on P14. Lower body weight could result from poor feeding due to increased sensitivity to the acute morphine physiological effects and/or morphine-induced withdrawal-induced gastrointestinal disruptions, contributing to internal distress.

Our results provide a comprehensive, in-depth investigation of morphine withdrawal-induced USV features in neonatal mice. We expanded on our previous findings of altered syllable profiles associated with morphine withdrawal using a large dataset across four FVB/N substrains. We conducted a more extensive analysis of the spectrotemporal properties of USVs and found withdrawal-induced changes in USV length, power, principal frequency, change in frequency, and peak frequency on P7 and P14. Additionally, we observed a less random syllable repertoire yet more unique syllable sequences in morphine-withdrawn pups compared to saline-treated pups on P14. Most sequences were emitted infrequently; thus, the syllable profile may communicate more important information than their temporal organization. We did not observe significant correlations between Complex 3 emissions and nociception phenotypes, suggesting that increased Complex 3 emissions is not associated with somatic withdrawal severity (at least not with stimulus-evoked nociception) but could serve as a biobehavioral marker for other withdrawal traits, including the negative affective state.

Importantly, our USV findings in a mouse model for NOWS align with recent human studies regarding withdrawal-induced alterations in cry features [33–36], suggesting that measuring more detailed cry properties using unbiased, objective methods may provide a more accurate assessment of NOWS symptom severity compared to solely subjective scoring systems currently used in clinical settings.

## Author Contributions

K.K.W.: Data collection, analysis, and writing of manuscript; T.M.: Data collection; S.A.M.: Data collection; C.S.M.: Data collection; K.J.: Data collection; N.M.A.; Data collection; K.T.R.: Data collection; M.B.R.; Data collection; J.A.B.; Data collection; E.M.W.: Writing of manuscript; C.D.B.: Experimental design, writing of manuscript.

