## Supplemental Figure Legends for "The acoustic properties, syllable structure, and syllable sequences of ultrasonic vocalizations (USVs) during neonatal opioid withdrawal in FVB/N mouse substrains"

**Figure S1. Morphine Treatment Interactions on USV properties during spontaneous opioid withdrawal on P7**. Data are plotted as the mean ± SEM. Data were analyzed using linear mixed models with Saline Treatment, Female Sex, and NCrl Substrain as the reference variables. **A.** **P7 Total USVs:** There was a Substrain x Morphine Treatment x Sex interaction (β = -190.01, SE = 76.74, t(167) = -2.48, **p* = 0.014) where morphine-withdrawn NTac females emitted more USVs than saline control NTac females (β = 278.03, SE = 115.28, t(159) = 2.41, **p* = 0.017). NTac: SAL, n = 16 (9F, 7M); MOR, n = 21 (12F, 9M). **B. P7 % Chevron:** There was a Substrain x Morphine Treatment x Sex interaction (β = 0.087, SE = 0.038, t(167) = 2.31, **p* = 0.022) where morphine-withdrawn NTac males emitted a greater percentage of Chevron syllables than saline control NTac males (β = 0.24, SE = 0.059, t(159) = 4.15, *****p* < 0.0001). Morphine-withdrawn NJ females emitted a greater percentage of Chevron syllables than saline control NJ females (β = 0.13, SE = 0.047, t(159) = 2.71, ***p* = 0.075). NTac: SAL, n = 16 (9F, 7M); MOR, n = 20 (11F, 9M). NJ: SAL, n = 25 (12F, 13M); MOR, n = 23 (13F, 10M). **C. P7 % Complex 4:** There was a Morphine Treatment x Sex interaction (β = 0.044, SE = 0.021, t(167) = 2.08, **p* = 0.039) where morphine-withdrawn males emitted a greater proportion of Complex 4 syllables than saline control males (β = 0.013, SE = 0.0062, t(167) = 2.08, **p* = 0.039). SAL, n = 88 (50F, 38M); MOR, n = 87 (45F, 42M). **D. P7 % Reverse Chevron:** There was a Substrain x Morphine Treatment x Sex interaction (β = -0.0062, SE = 0.0072, t(167) = -2.29, **p* = 0.023) where morphine-withdrawn NTac females emitted a greater proportion of Reverse Chevron syllables compared to saline control NTac females (β = 0.014, SE = 0.0041, t(159) = 3.35, ***p* = 0.0010). NTac: SAL, n = 16 (9F, 7M); MOR, n = 20 (11F, 9M). **E.** **P7 Power:** There was a Substrain x Morphine Treatment x Sex interaction (β = -2.83, SE = 1.41, t(131) = -2.01, **p* = 0.047) where morphine-withdrawn NCrl females emitted quieter USVs (β = -4.86, SE = 1.92, t(123) = -2.53, **p* = 0.013) than saline-control NCrl females. Additionally, morphine-withdrawn NTac males emitted quieter USVs (β = -7.74, SE = 2.22, t(123) = -3.49, ****p* = 0.0007) than saline-control NTac males. NCrl: SAL, n = 19 (13F, 6M); MOR, n = 16 (8F, 8M). NTac: SAL, n = 14 (7F, 7M); MOR = 18 (10F, 8M).

**Figure S2. Morphine Treatment Interactions on USV syllables during spontaneous opioid withdrawal on P14.** Data are plotted as the mean ± SEM. Data were analyzed using linear mixed models with Saline Treatment, Female Sex, and NCrl Substrain as the reference variables. **A. P14 % Downward:** There was a Substrain x Morphine Treatment x Sex interaction (β = 0.042, SE = 0.014, t(144) = 2.98, ***p* = 0.0034) where morphine-withdrawn NTac females emitted fewer Downward syllables compared to saline control NTac females (β = -0.051, SE = 0.022, t(136) = 2.30, **p* = 0.023). NTac: SAL, n = 17 (9F, 8M); MOR, n = 18 (10F, 8M). **B. P14 % Upward:** There was a Substrain x Morphine Treatment x Sex interaction (β = -0.059, SE = 0.027, t(144) = -2.21, **p* = 0.029) where morphine-withdrawn NTac males emitted fewer Upward syllables compared to saline control NTac males (β = -0.12, SE = 0.041, t(136) = -2.98, ***p* = 0.0034), morphine-withdrawn NCrl females emitted fewer Upward syllables compared to saline control NCrl females (β = -0.098, SE = 0.034, t(136) = -2.89, ***p* = 0.0045), morphine-withdrawn NHsd females emitted fewer Upward syllables compared to saline control NHsd females (β = -0.087, SE = 0.035, t(136) = -2.52, **p* = 0.013), and morphine-withdrawn NJ females emitted fewer Upward syllables compared to saline control NJ females (β = 0.097, SE = 0.039, t(136) = -2.46, **p* = 0.015). NTac: SAL, n = 17 (9F, 8M); MOR, n = 18 (10F, 8M). NCrl: SAL, n = 20 (12F, 8M); MOR, n = 20 (11F, 9M). NHsd: SAL, n = 21 (12F, 9M); MOR, n = 19 (10F, 9M). NJ: SAL, n = 20 (8F, 12M); MOR, n = 17 (9F, 8M).

**Figure S3. Morphine Treatment Interactions on P7 USV syllable features during spontaneous opioid withdrawal.** Data are plotted as the mean ± SEM. Data were analyzed using linear mixed models with Saline Treatment, Female Sex, and NCrl Substrain as the reference variables. **A. P7 Downward Low Frequency**: There was a Substrain x Morphine Treatment x Sex interaction (β = 5.47, SE = 2.44, t(124) = 2.24, **p* = 0.029) where morphine-withdrawn NTac males emitted a higher low frequency than saline control NTac males (β = 9.34, SE = 4.19, t(116) = 2.23, **p* = 0.028) and morphine-withdrawn NCrl females emitted a higher low frequency than saline control NCrl females (β = 6.59, SE = 3.26, t(116) = 2.02, **p* = 0.046). **B. P7 High Frequency**: There was a Substrain x Morphine Treatment x Sex interaction (β = 10.11, SE = 3.06, t(124) = 3.31, ***p* = 0.0012) where morphine-withdrawn NTac males emitted higher frequency Downward syllables than saline control NTac males (β = 15.43, SE = 5.18, t(116) = 2.98, ***p* = 0.0036) and morphine-withdrawn NCrl males emitted lower high frequency Downward syllables than saline control NCrl males (β = -10.88, SE = 4.82, t(116) = 2.26, **p* = 0.026). **C. P7 Downward Change in Frequency**: There was a Substrain x Morphine Treatment x Sex interaction (β = 4.64, SE = 1.48, t(124) = 3.14, ***p* = 0.0021) where morphine-withdrawn NTac males emitted a greater change in frequency of Downward syllables than saline control NTac males (β = 15.42, SE = 5.18, t(116) = 2.98, **p* = 0.012) and morphine-withdrawn NCrl males emitted a reduced change in frequency of Downward syllables than saline control NCrl males (β = -5.50, SE = 2.23, t(116) = -2.46, **p* = 0.015). **D. P7 Downward Principal Frequency**: There was a Substrain x Morphine Treatment x Sex interaction (β = 8.32, SE = 2.83, t(124) = 2.94, ***p* = 0.0039) where morphine-withdrawn NTac males emitted a higher principal frequency than saline control NTac males (β = 15.14, SE = 4.86, t(116) = 3.11, ***p* = 0.0023). **E. P7 Downward Peak Frequency:** There was a Substrain x Morphine Treatment x Sex interaction (β = 8.46, SE = 2.69, t(124) = 3.13, ***p* = 0.0022) where morphine-withdrawn NTac males emitted a higher peak frequency of Downward syllables than saline control NTac males (β = 15.29, SE = 4.58, t(116) = 3.34, ***p* = 0.0011) and morphine-withdrawn NCrl females emitted a higher peak frequency of Downward syllables than saline control NCrl females (β = 7.50, SE = 3.57, t(116) = 2.10, **p* = 0.038). NTac: SAL, n = 11 (5F, 6M); MOR, n = 14 (7F, 7M). NCrl: SAL, n = 20 (13F, 7M); MOR, n = 17 (9F, 8M). **F. P7 Complex 4 Power:** There was a Morphine Treatment x Sex interaction (β = 7.37, SE = 3.13, t(104) = 2.35, **p* = 0.020) where morphine-withdraw NCrl pups emitted reduced power compared to saline control pups (β = -7.55 SE = 2.07, t(96) = -3.66, ****p* = 0.00042). SAL, n = 16 (9F, 7M); MOR, n = 11 (7F, 4M). **G. P7 Flat Frequencies**: High Frequency: There was a Morphine Treatment x Sex interaction (β = 3.95, SE = 2.03, t(129) = 1.94, **p* = 0.054) where morphine-withdrawn females emitted a greater high frequency of Flat syllables compared to saline control females (β = 3.35, SE = 1.50, t(121) = 2.23, **p* = 0.028). Principal Frequency: There was a Morphine Treatment x Sex interaction (β = -11.65, SE = 5.48, t(129) = -2.13, **p* = 0.036) where morphine-withdrawn females emitted a greater principal frequency of Flat syllables compared to saline control females (β = 3.96, SE = 1.54, t(121) = 2.57, **p* = 0.011). Peak Frequency: There was a Morphine Treatment x Sex interaction (β = -11.59, SE = 5.45, t(129) = -2.13, **p* = 0.035) where morphine-withdrawn females emitted a greater peak frequency of Flat syllables compared to saline control females (β = 4.01, SE = 1.53, t(121) = 2.62, ***p* = 0.0098). SAL, n = 73 (40F, 33M); MOR, n = 64 (37F, 27M). **H. P7 Upward Length:** There was a Substrain x Morphine Treatment x Sex interaction (β = 8.31, SE = 3.05, t(130) = 2.73, ***p* = 0.0073) where morphine-withdrawn NTac females emitted shorter Upward syllables compared to saline control NTac females (β = -14.8, SE = 4.94, t(122) = -2.99, ***p* = 0.0033. NTac: SAL, n = 16 (6F, 10M); MOR, n = 16 (9F, 7M).

**Figure S4. Morphine treatment alters USV syllable acoustic features in FVB/N pups on P7 during spontaneous opioid withdrawal.** Data are plotted as the mean ± SEM. Data were analyzed using linear mixed models with Saline Treatment as the reference variable. The following data are collapsed across Substrain and Sex for simplicity.

| **Syllable** | **A. Length** | **B. Power** | **C. Low Frequency** | **D. High Frequency** | **E. Frequency Change** | **F. Principal Frequency** | **G. Peak Frequency** | **n** |
| --- | --- | --- | --- | --- | --- | --- | --- | --- |
| **Chevron** | (β = -0.0035, SE = 0.0018, t(143) = -2.01, **p* = 0.046) | (β = -2.33, SE = 0.68, t(143) = -3.43, ****p* = 0.00078) | (β = -0.11, SE = 0.92, t(143) = -0.12, *p* = 0.91) | (β = -1.39, SE = 1.15, t(143) = -1.21, *p* = 0.23) | (β = -1.28, SE = 0.57, t(143) = -2.26, **p* = 0.026) | (β = -0.11, SE = 1.08, t(143) = -0.11, *p* = 0.92) | (β = 0.88, SE = 1.00, t(143) = 0.88, *p* = 0.34) | SAL = 77 |
|  |  |  |  |  |  |  |  | MOR = 68 |
| **Complex** | (β = -0.0047, SE = 0.0024, t(141) = -1.99, **p* = 0.048) | (β = -2.44, SE = 0.80, t(141) = -3.04, ***p* = 0.0028) | (β = -1.38, SE = 1.16, t(141) = -1.91, *p* = 0.24) | (β = -2.07, SE = 1.46, t(141) = -1.42, *p* = 0.16) | (β = -0.69, SE = 0.80, t(141) = -0.87, *p* = 0.39) | (β = -0.97, SE = 1.34, t(141) = -0.73, *p* = 0.47) | (β = 0.67, SE = 1.26, t(141) = 0.52, *p* = 0.60) | SAL = 75 |
|  |  |  |  |  |  |  |  | MOR = 68 |
| **Complex 2** | (β = -0.0064, SE = 0.0026, t(141) = -2.48, **p* = 0.014) | (β = -3.12, SE = 0.80, t(141) = -3.91, ****p* = 0.000014) | (β = 2.82, SE = 0.85, t(141) = 3.31, ***p* = 0.0012) | (β = 3.33, SE = 1.43, t(141) = 2.33, **p* = 0.021) | (β = 0.51, SE = 1.22, t(141) = 0.42, *p* = 0.67) | (β = 4.62, SE = 1.29, t(141) = 3.58, ****p* = 0.00048) | (β = 5.66, SE = 1.25, t(141) = 4.54, *****p* < 0.0001) | SAL = 77 |
|  |  |  |  |  |  |  |  | MOR = 66 |
| **Complex 3** | (β = -0.0071, SE = 0.0020, t(142) = -3.51, ****p* = 0.00059) | (β = -3.66, SE = 0.73, t(142) = -5.01, *****p* < 0.0001) | (β = 2.80, SE = 0.48, t(142) = 5.78, *****p* < 0.0001) | (β = 1.62, SE = 1.22, t(142) = 1.32, *p* = 0.19) | (β = -1.18, SE = 1.12, t(142) = -1.06, *p* = 0.29) | (β = 3.99, SE = 0.69, t(142) = 5.78, *****p* < 0.0001) | (β = 3.97, SE = 0.73, t(142) = 5.40, *****p* < 0.0001) | SAL = 77 |
|  |  |  |  |  |  |  |  | MOR = 67 |
| **Complex 4** | (β = -0.0051, SE = 0.0032, t(116) = -1.58, *p* = 0.12) | (β = -3.43, SE = 1.00, t(116) = -3.42, ****p* = 0.00087) | (β = 4.21, SE = 1.58, t(116) = 2.66, ***p* = 0.0089) | (β = 1.64, SE = 1.59, t(116) = 1.03, *p* = 0.30) | (β = -2.57, SE = 2.09, t(116) = -1.23, *p* = 0.22) | (β = 2.96, SE = 1.21, t(116) = 2.44, **p* = 0.016) | (β = 2.92, SE = 1.27, t(116) = 2.30, **p* = 0.023) | SAL = 67 |
|  |  |  |  |  |  |  |  | MOR = 51 |
| **Downward** | (β = -0.0012, SE = 0.0029, t(137) = -0.42, *p* = 0.68) | (β = -1.84, SE = 0.83, t(137) = -2.21, **p* = 0.029) | (β = 2.01, SE = 1.24, t(137) = 1.50, *p* = 0.14) | (β = 1.08, SE = 1.72, t(137) = 0.63, *p* = 0.53) | (β = -0.92, SE = 0.84, t(137) = -1.10, *p* = 0.27) | (β = 2.45, SE = 1.57, t(137) = 1.56, *p* = 0.12) | (β = 2.68, SE = 1.50, t(137) = 1.79, *p* = 0.076) | SAL = 73 |
|  |  |  |  |  |  |  |  | MOR = 66 |
| **Flat** | (β = -0.0049, SE = 0.0018, t(142) = -2.73, ***p* = 0.0071) | (β = -3.08, SE = 0.83, t(142) = -3.68, ****p* = 0.00033) | (β = 2.85, SE = 1.07, t(142) = 2.67, ***p* = 0.0085) | (β = 2.55, SE = 1.09, t(142) = 2.34, **p* = 0.021) | (β = -0.30, SE = 0.34, t(142) = -0.88, *p* = 0.38) | (β = 3.00, SE = 1.09, t(142) = 2.74, ***p* = 0.0069) | (β = -3.05, SE = 1.09, t(142) = 2.81, ***p* = 0.0057) | SAL = 77 |
|  |  |  |  |  |  |  |  | MOR = 67 |
| **Reverse Chevron** | (β = -0.0052, SE = 0.0053, t(86) = -0.99, *p* = 0.33) | (β = -2.34, SE = 1.30, t(86) = -1.80, *p* = 0.076) | (β = 2.98, SE = 2.37, t(86) = 1.26, *p* = 0.21) | (β = 4.02, SE = 2.64, t(86) = 1.52, *p* = 0.13) | (β = 1.04, SE = 1.41, t(86) = 0.73, *p* = 0.47) | (β = 5.58, SE = 2.43, t(86) = 2.29, **p* = 0.024) | (β = 4.88, SE = 2.37, t(86) = 2.06, **p* = 0.043) | SAL = 47 |
|  |  |  |  |  |  |  |  | MOR = 41 |
| **Short** | (β = -0.00082, SE = 0.00018, t(139) = -0.47, *p* = 0.64) | (β = -0.40, SE = 0.71, t(139) = -0.56, *p* = 0.58) | (β = 2.41, SE = 0.87, t(139) = 2.78, ***p* = 0.0063) | (β = 2.58, SE = 0.90, t(139) = 2.87, ***p* = 0.0047) | (β = 0.17, SE = 0.62, t(139) = 0.27, *p* = 0.79) | (β = 2.55, SE = 0.84, t(139) = 3.03, ***p* = 0.0029) | (β = -0.40, SE = 0.71, t(139) = -0.56, *p* = 0.58) | SAL = 76 |
|  |  |  |  |  |  |  |  | MOR = 65 |
| **Upward** | (β = -0.00028, SE = 0.0017, t(143) = -0.17, *p* = 0.86) | (β = -2.88, SE = 0.75, t(143) = -3.84, ****p* = 0.00018) | (β = 1.70, SE = 0.78, t(143) = 2.16, **p* = 0.032) | (β = 2.18, SE = 0.76, t(143) = 2.86, ***p* = 0.0049) | (β = 0.48, SE = 0.44, t(143) = 1.11, *p* = 0.27) | (β = 1.84, SE = 0.78, t(143) = 2.34, **p* = 0.021) | (β = 2.38, SE = 0.75, t(143) = 3.18, ***p* = 0.0018) | SAL, n = 77 |
|  |  |  |  |  |  |  |  | MOR, n = 68 |

**Figure S5. Morphine Treatment Interactions on P14 USV syllable features during spontaneous opioid withdrawal.** Data are plotted as the mean ± SEM. Data were analyzed using linear mixed models with Saline Treatment, Female Sex, and NCrl Substrain as the reference variables. **A. P14 Chevron High Frequency:** There was a Morphine Treatment x Substrain interaction (β = 3.67, SE = 1.84, t(117) = 2.00, **p* = 0.048) where morphine-withdrawn NCrl pups emitted lower high frequency Chevron syllables compared to saline control NCrl pups (β = -4.41, SE = 1.43, t(109) = -3.09, ***p* = 0.0025) and morphine-withdrawn NTac pups emitted lower high frequency Chevron syllables than saline control NCrl pups (β = -2.90, SE = 1.12, t(109) = -2.60, **p* = 0.011). **B. P14 Chevron Principal Frequency**: There was a Morphine Treatment x Substrain interaction (β = 3.78, SE= 1.64, t(117) = 2.30, **p* = 0.023) where morphine-withdrawn NJ pups emitted higher principal frequency Chevron syllables compared to saline control NJ pups (β = 2.40, SE = 0.97, t(109) = 2.47, **p* = 0.015), morphine-withdrawn NCrl pups emitted lower principal frequency Chevron syllables compared to saline control NCrl pups (β = -3.09, SE = 1.27, t(109) = -2.44, **p* = 0.016), and morphine-withdrawn NTac pups emitted higher principal frequency Chevron syllables compared to saline control NTac pups (β = 2.38, SE = 1.18, t(109) = 2.03, **p* = 0.045). **C. P14 Chevron Peak Frequency:** There was a Morphine Treatment x Substrain interaction (β = 4.28, SE = 1.86, t(117) = 2.30, **p* = 0.023) where morphine-withdrawn NCrl pups emitted a lower peak frequency than saline control NCrl pups (β = -3.12, SE = 1.46, t(117) = -2.13, **p* = 0.035). NCrl: SAL, n = 17 (11F, 6M); MOR, n = 16 (6F, 10M). **D. P14 Complex Power:** There was a Morphine Treatment x Sex interaction (β = -17.21, SE = 7.87, t(95) = -2.19, **p* = 0.031) where morphine-withdrawn males emit reduced power than saline control males (β = -4.91, SE = 2.18, t(87) = -2.25, **p* = 0.027). SAL, n = 47 (30F, 17M); MOR, n = 56 (25F, 31M). **E. P14 Complex 4 Low Frequency**: There was a Morphine Treatment x Substrain interaction (β = -5.99, SE = 2.89, t(77) = -2.08, **p* = 0.041) where morphine-withdrawn NTac pups emitted reduced low frequency compared to saline control NTac pups (β = -4.03, SE = 1.85, t(69) = -2.18, **p* = 0.033). **F. P14 Reverse Chevron Length:** There was a Morphine Treatment x Sex interaction (β = -23.79, SE = 4.24, t(87) = -2.04, **p* = 0.045) where morphine-withdrawn females emitted longer Reverse Chevron syllables than saline control females (β = 7.78, SE = 2.75, t(79) = 2.83, ***p* = 0.059). SAL, n = 42 (27F, 15M); MOR, n = 53 (25F, 28M). **G. P14 Upward Peak Frequency:** There was a Substrain x Morphine Treatment x Sex interaction (β = -2.45, SE = 1.22, t(122) = -2.01, **p* = 0.047) where morphine-withdrawn NJ males emitted higher peak frequencies than saline control NJ males (β = 4.09, SE = 1.58, t(114) = 2.60, **p* = 0.011) and morphine-withdrawn NHsd females emitted lower peak frequencies than saline control NHsd females (β = -3.64, SE = 1.64, t(114) = -2.22, **p* = 0.029). NJ: SAL, n = 20 (8F, 12M); MOR, n = 16 (8F, 8M). NHsd: SAL, n = 20 (12F, 8M); MOR, n = 16 (7F, 9M).

**Figure S6. Morphine Treatment Interactions on P14 Complex 3 features during spontaneous opioid withdrawal.** Data are plotted as the mean ± SEM. Data were analyzed using linear mixed models with Saline Treatment, Female Sex, and NCrl Substrain as the reference variables**. A. P14 Complex 3 Low Frequency:** There was a Substrain x Morphine Treatment x Sex interaction (β = -3.09, SE = 0.96, t(118) = -3.22, ***p* = 0.0016) where morphine-withdrawn NCrl males emitted higher low frequency Complex 3 syllables than saline control NCrl males (β = 4.98, SE = 1.66, t(110) = 2.99, ***p* = 0.0034), morphine-withdrawn NHsd males emitted higher low frequency Complex 3 syllables than saline control NHsd males (β = 2.79, SE = 1.25, t(110) = 2.23, **p* = 0.028), and morphine-withdrawn NJ males emitted higher low frequency Complex 3 syllables than saline control NJ males (β = 2.47, SE = 1.18, t(110) = 2.10, **p* = 0.038). **B. P14 Complex 3 High Frequency:** There was a Substrain x Morphine Treatment interaction (β = 6.54, SE = 3.19, t(118) = 2.05, **p* = 0.043) where morphine-withdrawn NHsd pups emitted lower high frequency Complex 3 syllables than saline control NHsd pups (β = -5.28 SE = 1.79, t(110 = 2.95, ***p* = 0.0039). **C. P14 Complex 3 Principal Frequency:** There was a Substrain x Morphine Treatment x Sex interaction (β = -4.86, SE = 2.06, t(118) = -2.36, **p* = 0.020) where morphine-withdrawn NCrl males emitted higher principal frequency Complex 3 syllables than saline control NCrl males (β = 9.08, SE = 3.68, t(110) = 2.47, **p* = 0.015). NCrl: SAL, n = 15 (11F, 4M); MOR, n = 9 (3F, 6M). NHsd: SAL, n = 20 (12F, 8M); MOR, n = 16 (7F, 9M). NJ: SAL, n = 20 (8F, 12M); MOR, n = 17 (9F, 8M).

**Figure S7. Morphine treatment alters USV syllable acoustic features in FVB/N pups on P14 during spontaneous opioid withdrawal.** Data are plotted as the mean ± SEM. Data were analyzed using linear mixed models with Saline Treatment as the reference variable. The following data are collapsed across Substrain and Sex for simplicity. **A. P14 Length:** Morphine-withdrawn pups emitted longer Reverse Chevron syllables compared to saline control pups (β = 4.57, SE = 1.95, t(95) = 2.34, **p* = 0.021). **B. P14 Power:** Morphine-withdrawn pups emitted higher power Reverse Chevron syllables compared to saline control pups (β = 2.97, SE = 1.39, t(95) = 2.13, **p* = 0.036). **C. P14 Low Frequency:** Morphine-withdrawn pups emitted higher low frequency Complex 3 syllables compared to saline control pups (β = 1.06, SE = 0.50, t(128) = 2.14, **p* = 0.034). **D. P14 High Frequency:** Morphine-withdrawn pups emitted lower high frequency Complex 3 syllables compared to saline control pups (β = 2.29, SE = 1.05, t(128) = 2.19, **p* = 0.030). **E. P14 Frequency Change:** Morphine-withdrawn pups emitted Complex 3 syllables with a smaller frequency change compared to saline control pups (β = -3.35, SE = 0.94, t(128) = -3.58, ****p* = 0.00048) and Downward syllables with a smaller frequency change compared to saline control pups (β = -1.89, SE = 0.96, t(112) = -1.99, **p* = 0.049). Reverse Chevron: SAL, n = 47; MOR, n = 41. Complex 3: SAL, n = 77; MOR, n = 67. Downward: SAL, n = 77; MOR, n = 63.

**Figure S8. Correlation of withdrawal traits in saline-treated FVB/N pups on P7 and P14**. Colors indicate Pearson’s correlation coefficient: blue = positive, red = negative. Darker colors reflect stronger correlations. **p* < 0.01, ***p* < 0.001, ****p* < 0.0001. **A. P7**: SAL, n = 49 (25F, 24M). **B. P14**: SAL, n = 46 (22F, 24M).
