## Supplemental Tables for "The acoustic properties, syllable structure, and syllable sequences of ultrasonic vocalizations (USVs) during neonatal opioid withdrawal in FVB/N mouse substrains"

**Table S1. Factor analysis of saline-treated pups on P7.**

| **P7 Saline** | | | | | |
| --- | --- | --- | --- | --- | --- |
|  | **MR1** | **MR2** | **MR3** | **MR4** | **MR5** |
| P7 Body Weight |  |  |  |  |  |
| P7 Body Temperature |  |  |  |  |  |
| Hot Plate Latency |  |  |  |  |  |
| Hot Plate Velocity |  |  |  |  |  |
| Tail Withdrawal Latency |  |  |  |  |  |
| Average USV Velocity |  |  |  |  |  |
| Total USV Distance |  |  |  | 0.707 |  |
| Total # USVs |  |  |  | 0.618 |  |
| Length |  |  |  |  |  |
| Principal Freq | 0.976 |  |  |  |  |
| Low Freq | 0.953 |  |  |  |  |
| High Freq | 0.777 |  | 0.621 |  |  |
| Freq Change |  |  | 0.962 |  |  |
| Peak Freq | 0.954 |  |  |  |  |
| Power |  |  |  |  |  |
| Chevron | 0.667 |  |  |  |  |
| Complex |  |  |  |  | 0.800 |
| Complex 2 | -0.506 | 0.668 |  |  |  |
| Complex 3 |  | -0.768 |  |  |  |
| Complex 4 |  |  |  |  |  |
| Downward |  | 0.646 |  |  |  |
| Flat |  |  |  |  |  |
| Reverse Chevron |  |  |  |  | 0.752 |
| Short |  |  |  |  |  |
| Upward |  |  |  |  |  |

**Table S2. Factor analysis of morphine-withdrawn pups on P7.**

| **P7 Morphine** | | | | | |
| --- | --- | --- | --- | --- | --- |
|  | **MR1** | **MR2** | **MR3** | **MR4** | **MR5** |
| P7 Body Weight |  |  |  | 0.885 |  |
| P7 Body Temperature |  |  |  |  |  |
| Hot Plate Latency |  |  |  |  |  |
| Hot Plate Velocity |  |  |  |  |  |
| Tail Withdrawal Latency |  |  |  |  | -0.551 |
| Average USV Velocity |  |  |  |  |  |
| Total USV Distance |  |  |  |  |  |
| Total # USVs |  |  |  |  |  |
| Length |  | 0.794 |  |  |  |
| Principal Freq | 0.996 |  |  |  |  |
| Low Freq | 0.943 |  |  |  |  |
| High Freq | 0.814 |  |  |  |  |
| Freq Change |  | 0.833 |  |  |  |
| Peak Freq | 0.967 |  |  |  |  |
| Power |  | 0.633 |  |  |  |
| Chevron | 0.753 |  |  |  |  |
| Complex |  |  |  |  | -0.642 |
| Complex 2 |  |  |  |  |  |
| Complex 3 |  | 0.673 |  |  |  |
| Complex 4 |  |  |  |  |  |
| Downward |  | -0.770 |  |  |  |
| Flat |  |  |  |  |  |
| Reverse Chevron |  |  | 0.708 |  |  |
| Short |  |  |  |  |  |
| Upward |  |  | 0.723 |  |  |

**Table S3. Factor analysis of saline-treated pups on P14.**

| **P14 Saline** | | | | | |
| --- | --- | --- | --- | --- | --- |
|  | **MR1** | **MR2** | **MR3** | **MR4** | **MR5** |
| P14 Body Weight |  |  |  |  |  |
| P14 Body Temperature |  |  |  |  | 0.506 |
| Hot Plate Latency |  |  |  |  | 0.630 |
| Hot Plate Velocity |  |  | -0.619 |  |  |
| Tail Withdrawal Latency |  |  |  |  |  |
| Average USV Velocity |  |  |  |  |  |
| Total USV Distance |  |  |  |  |  |
| Total # USVs |  |  |  |  |  |
| Length |  | 0.637 | 0.679 |  |  |
| Principal Freq | 0.969 |  |  |  |  |
| Low Freq | 0.815 |  |  |  |  |
| High Freq | 0.783 | 0.572 |  |  |  |
| Freq Change |  | 0.952 |  |  |  |
| Peak Freq | 0.918 |  |  |  |  |
| Power |  |  | 0.629 |  |  |
| Chevron |  |  | 0.504 | -0.502 |  |
| Complex |  |  |  |  |  |
| Complex 2 |  |  |  | 0.622 |  |
| Complex 3 |  | 0.961 |  |  |  |
| Complex 4 |  |  |  |  |  |
| Downward | -0.540 |  |  |  |  |
| Flat |  |  |  |  |  |
| Reverse Chevron |  |  |  |  |  |
| Short |  |  | -0.717 |  |  |
| Upward |  | -0.624 |  |  |  |

**Table S4. Factor analysis of morphine-withdrawn pups on P14.**

| **P14 Morphine** | | | | | |
| --- | --- | --- | --- | --- | --- |
|  | **MR1** | **MR2** | **MR3** | **MR4** | **MR5** |
| P14 Body Weight |  | -0.535 |  |  |  |
| P14 Body Temperature |  |  |  |  |  |
| Hot Plate Latency |  |  |  | 0.864 |  |
| Hot Plate Velocity |  |  |  |  |  |
| Tail Withdrawal Latency |  |  |  | 0.577 |  |
| Average USV Velocity |  |  |  |  |  |
| Total USV Distance |  |  |  |  |  |
| Total # USVs |  |  |  |  |  |
| Length |  | 0.573 |  |  | 0.652 |
| Principal Freq | 0.952 |  |  |  |  |
| Low Freq | 0.930 |  |  |  |  |
| High Freq |  | 0.873 |  |  |  |
| Freq Change |  | 0.915 |  |  |  |
| Peak Freq | 0.907 |  |  |  |  |
| Power |  |  |  |  | 0.738 |
| Chevron |  |  | 0.840 |  |  |
| Complex |  |  |  |  |  |
| Complex 2 |  |  | -0.600 | 6.0 |  |
| Complex 3 | -0.561 | 0.705 |  |  |  |
| Complex 4 |  |  |  |  |  |
| Downward |  | -0.702 |  |  |  |
| Flat |  |  |  |  |  |
| Reverse Chevron |  |  |  |  |  |
| Short |  |  |  |  | -0.635 |
| Upward | 0.580 |  |  |  |  |
