## Supplementary figures and images for "The acoustic properties, syllable structure, and syllable sequences of ultrasonic vocalizations (USVs) during neonatal opioid withdrawal in FVB/N mouse substrains"

### Supplemental Figures

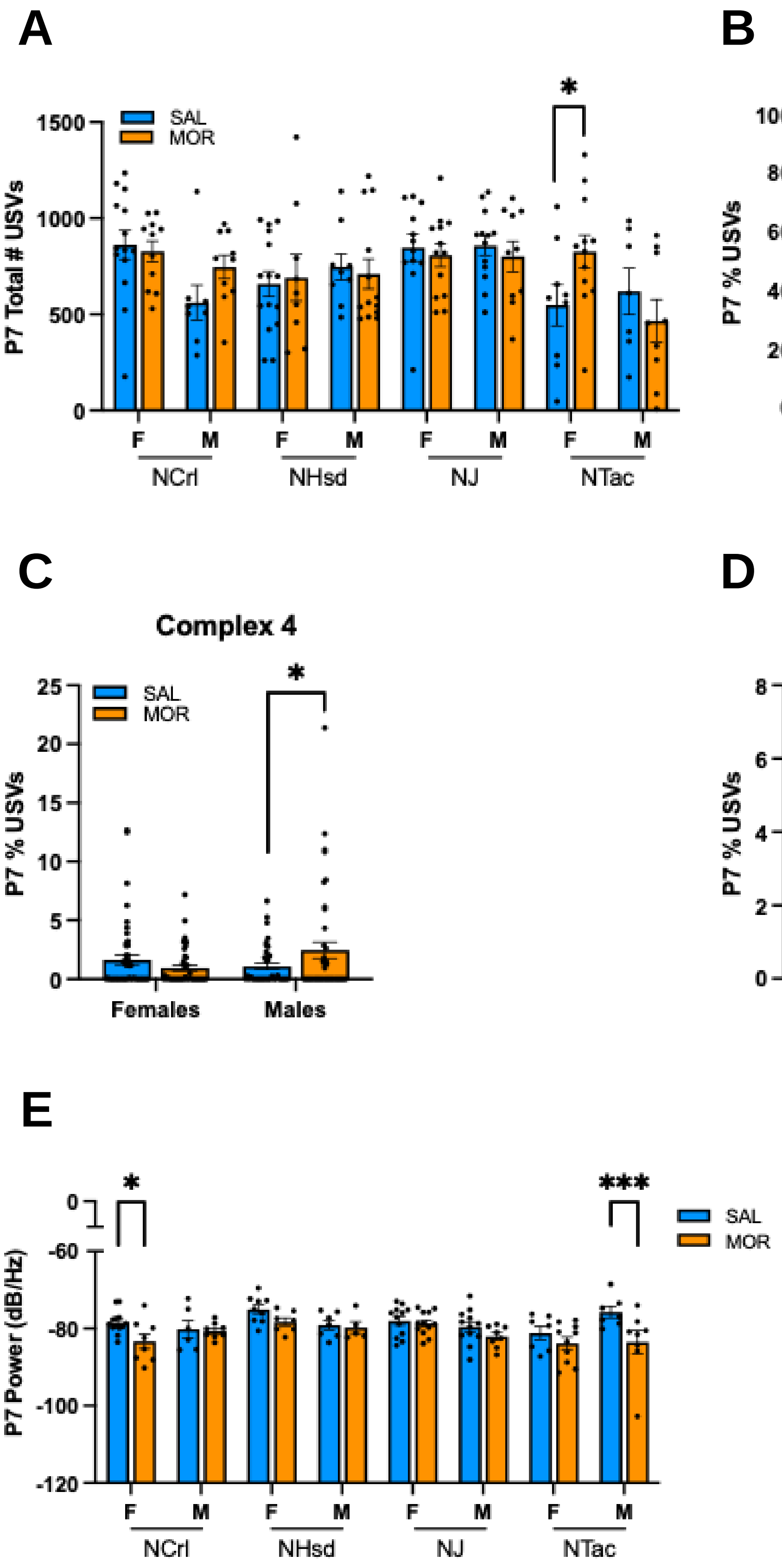

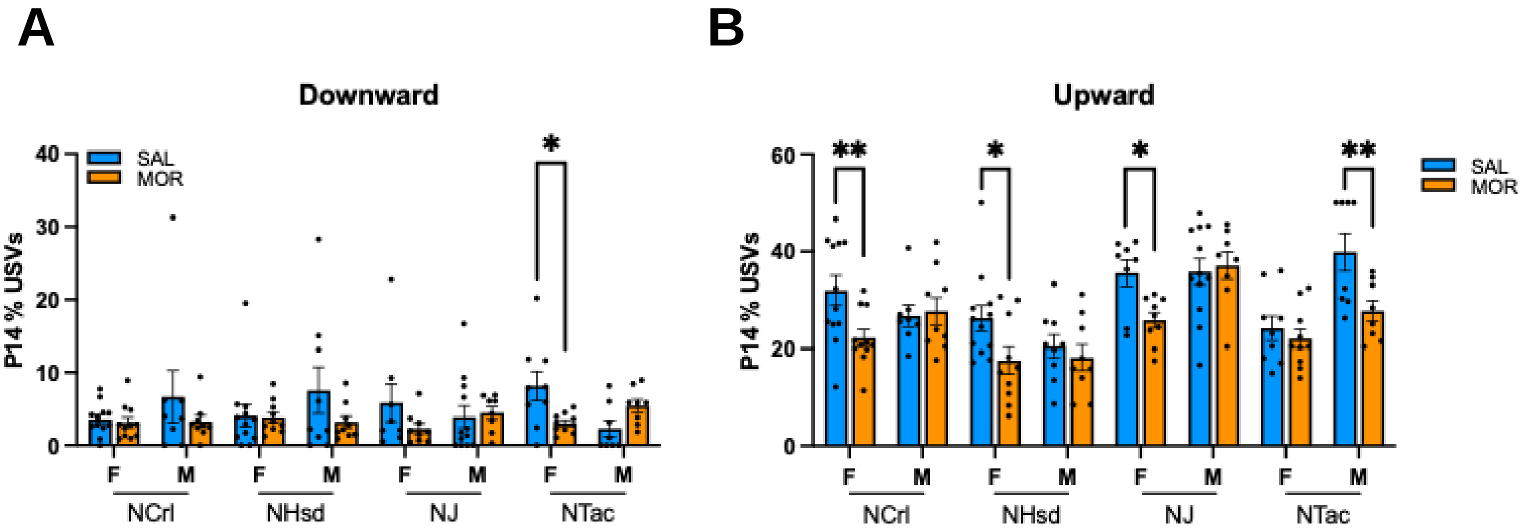

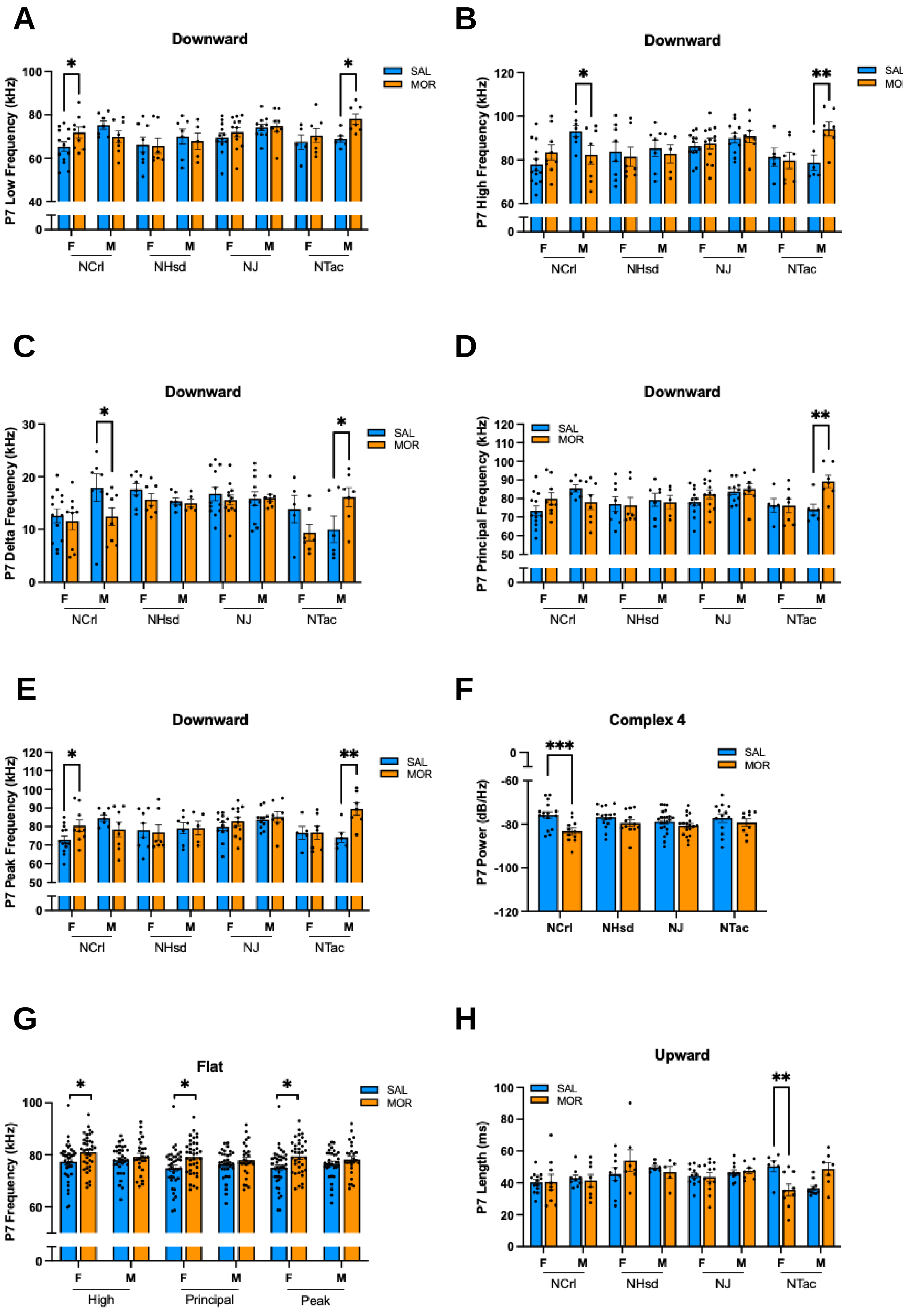

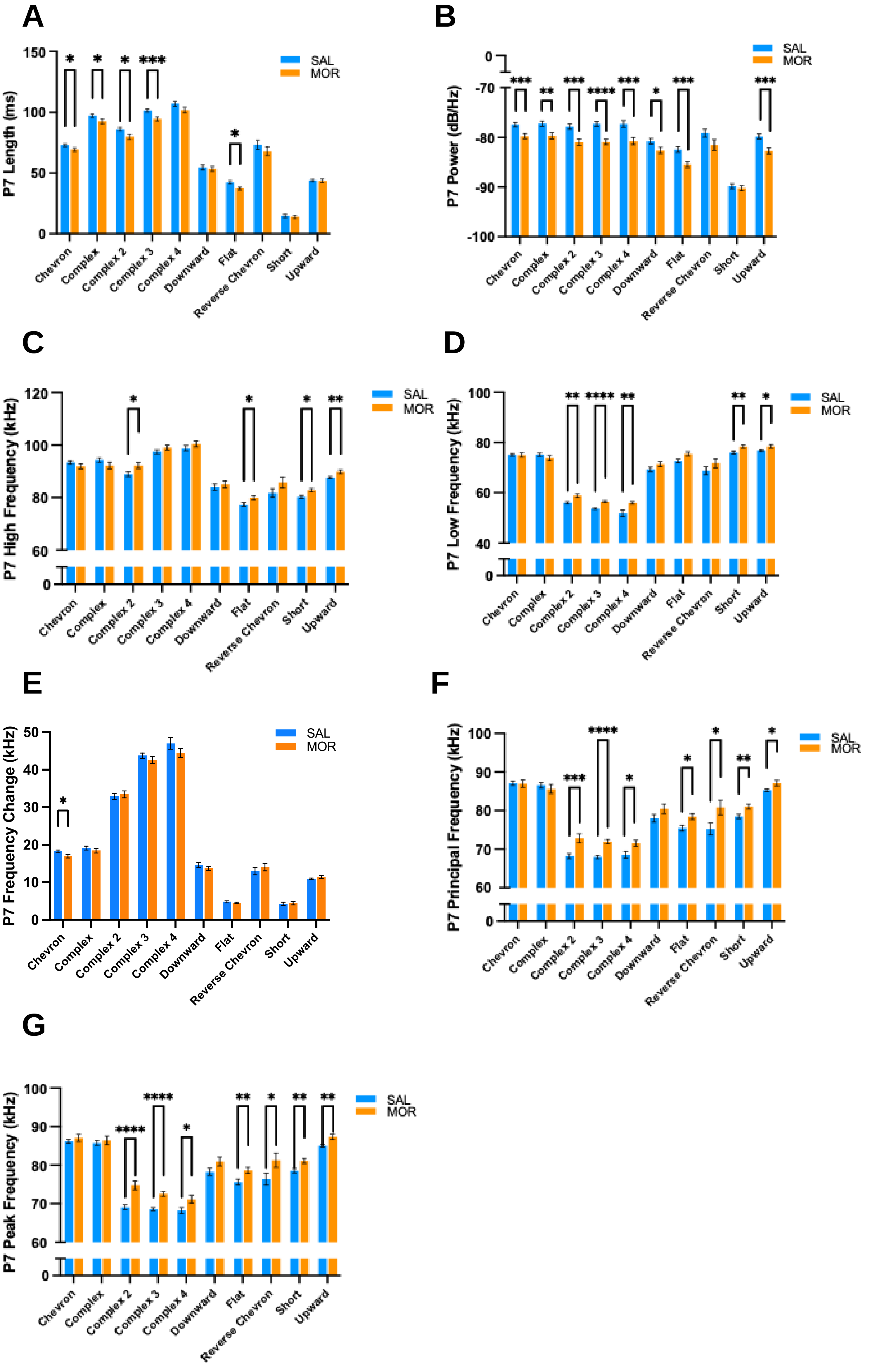

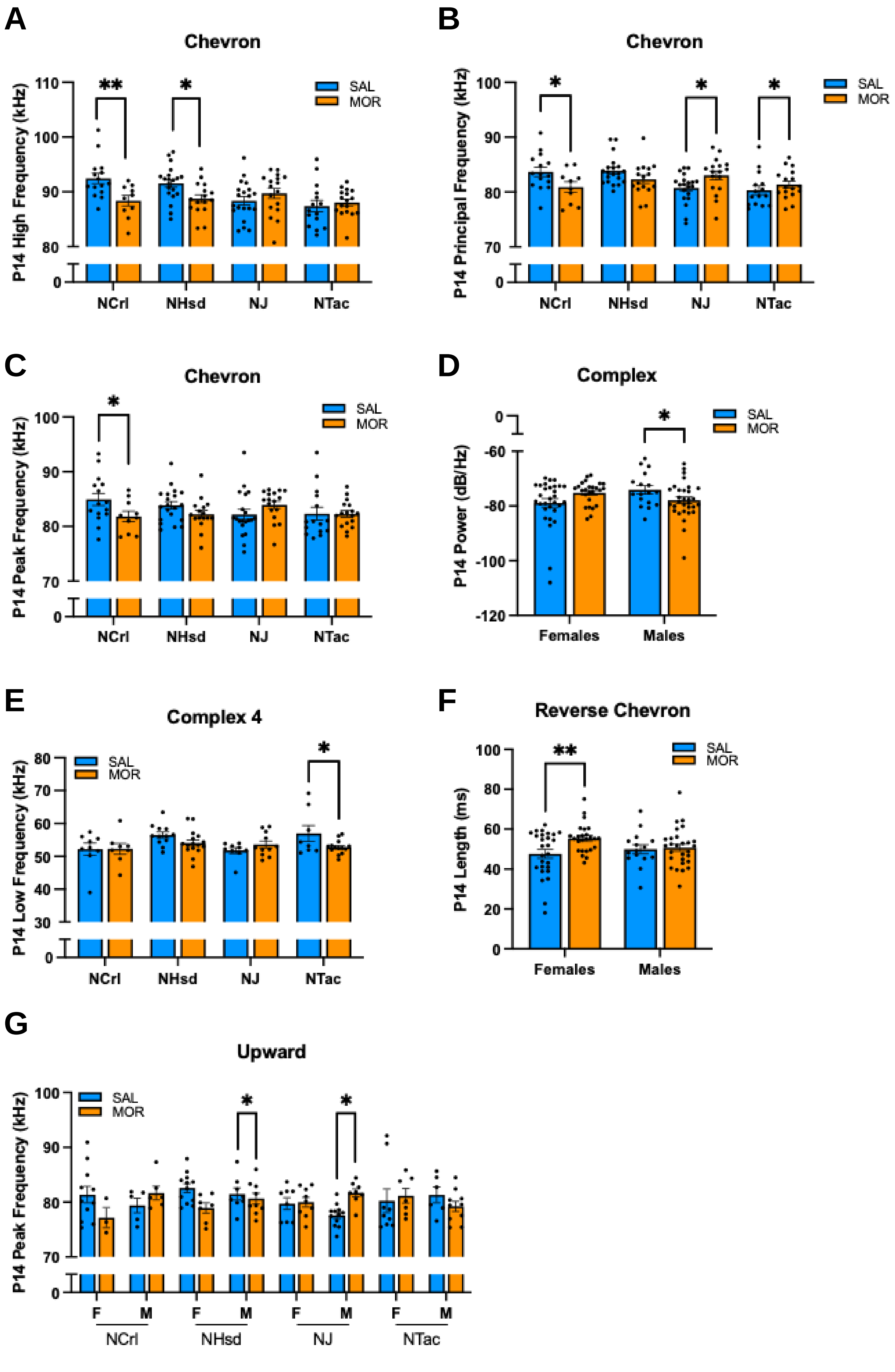

#### Figure S6

A

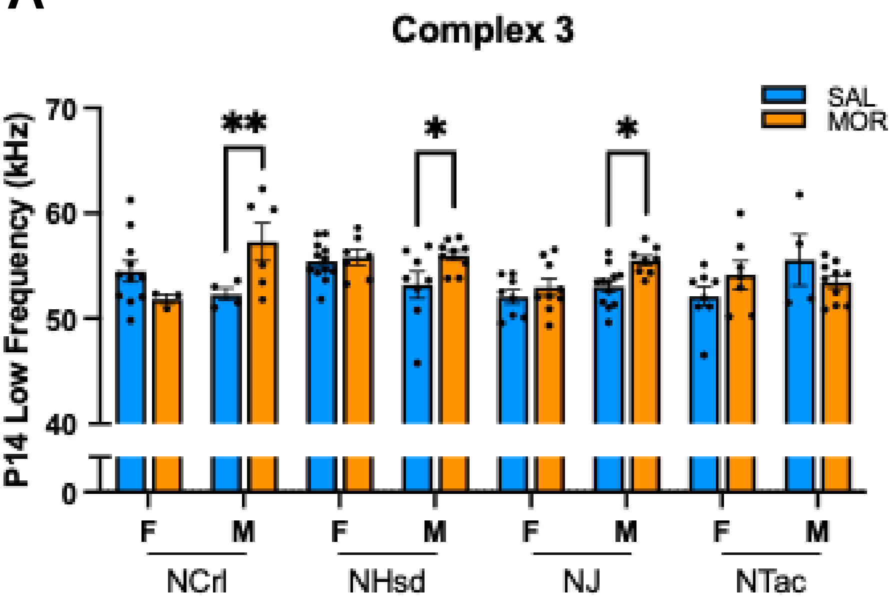


B

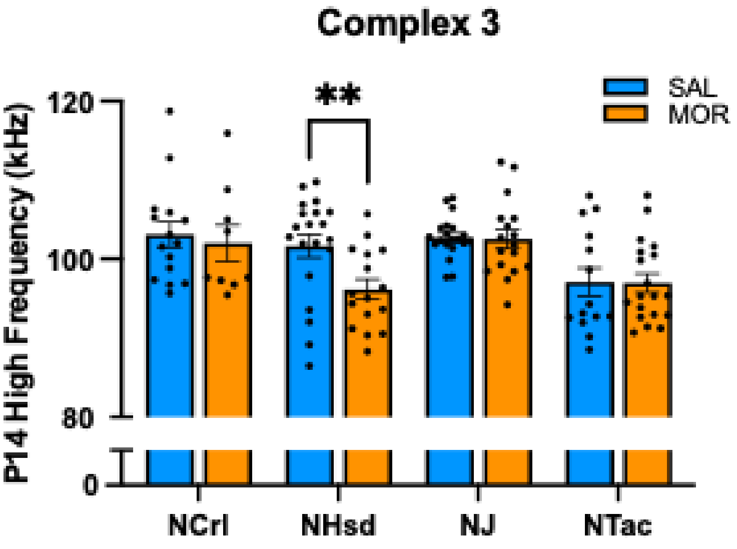


C

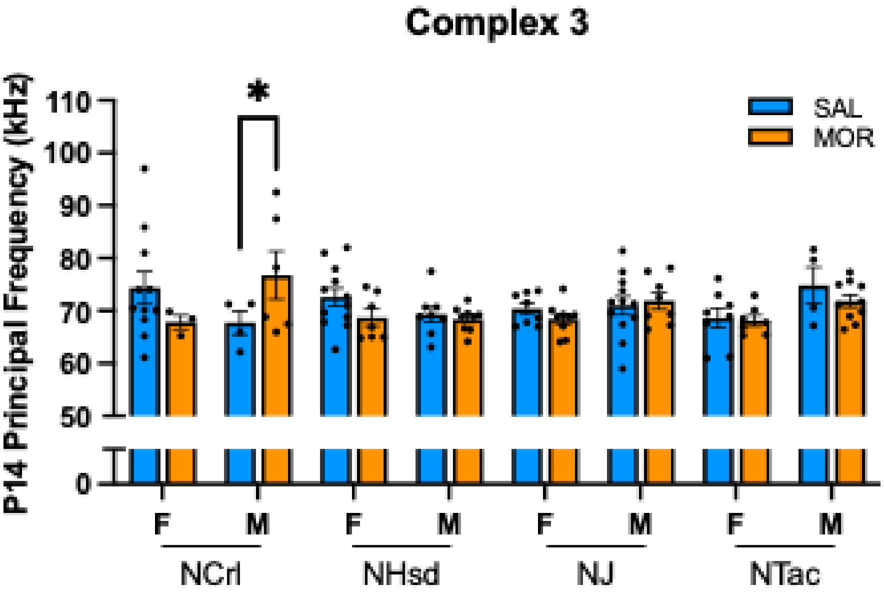

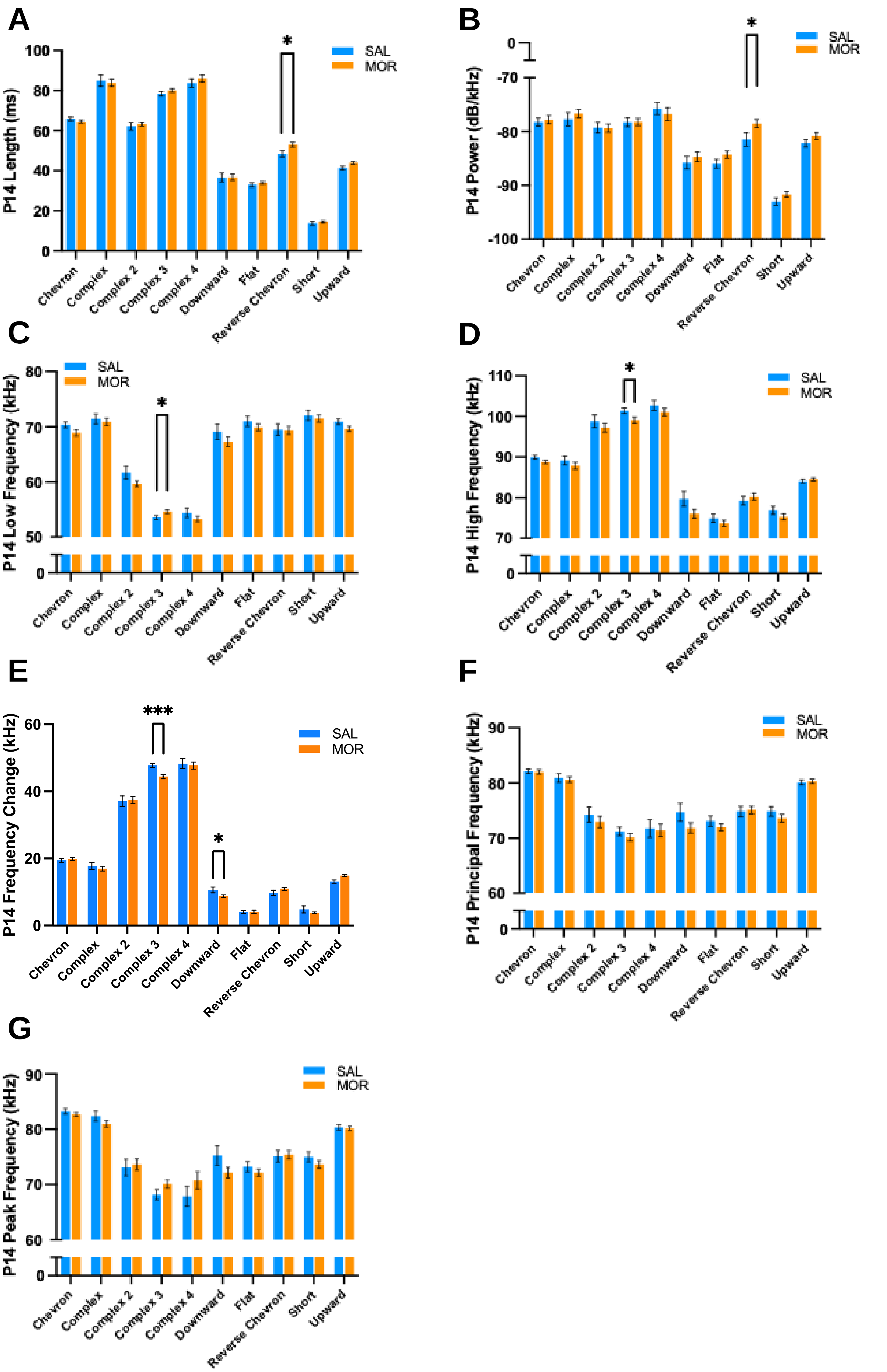

#### Figure S8

**A**

##### P7 Saline

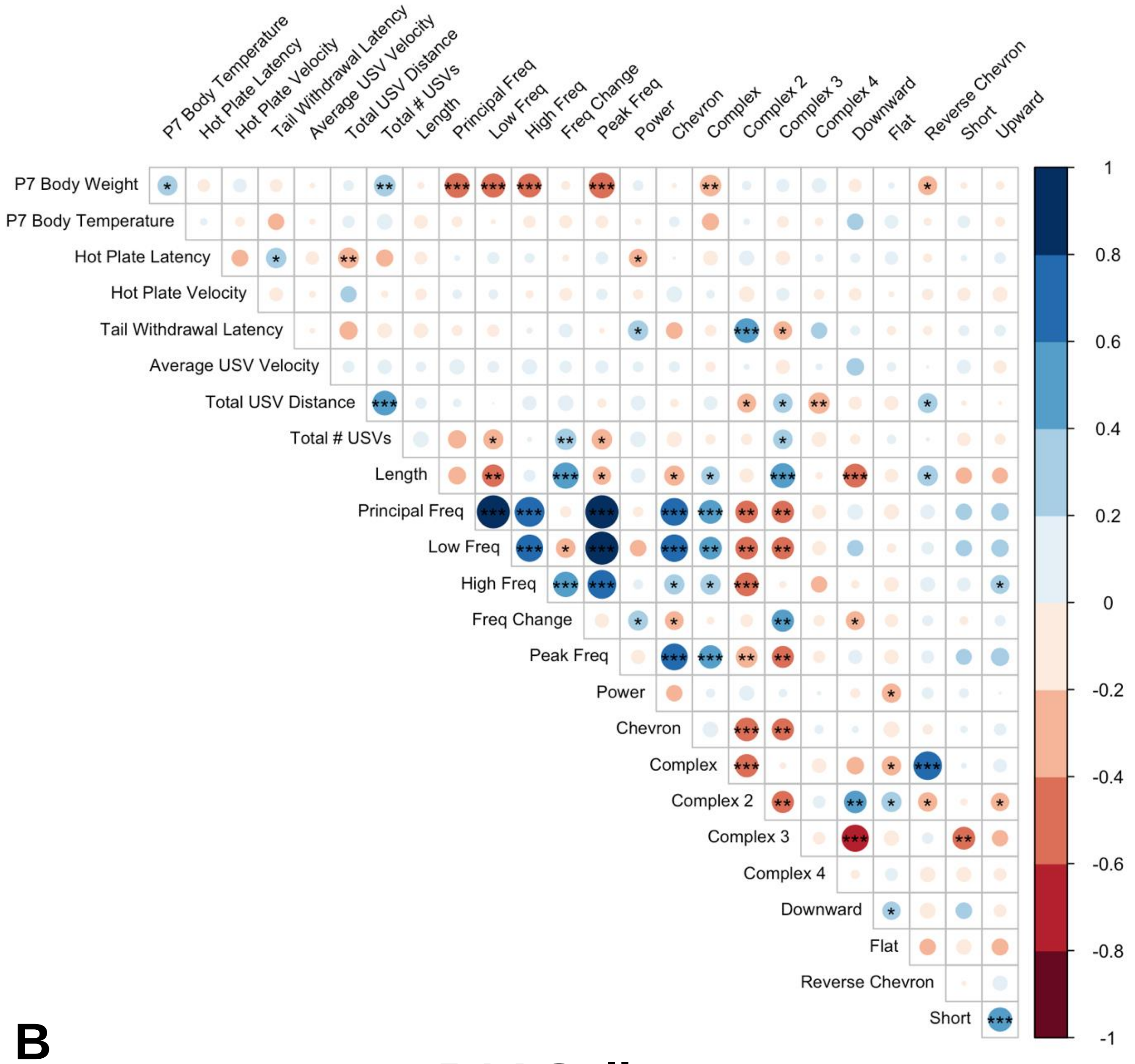


**B**

##### P14 Saline

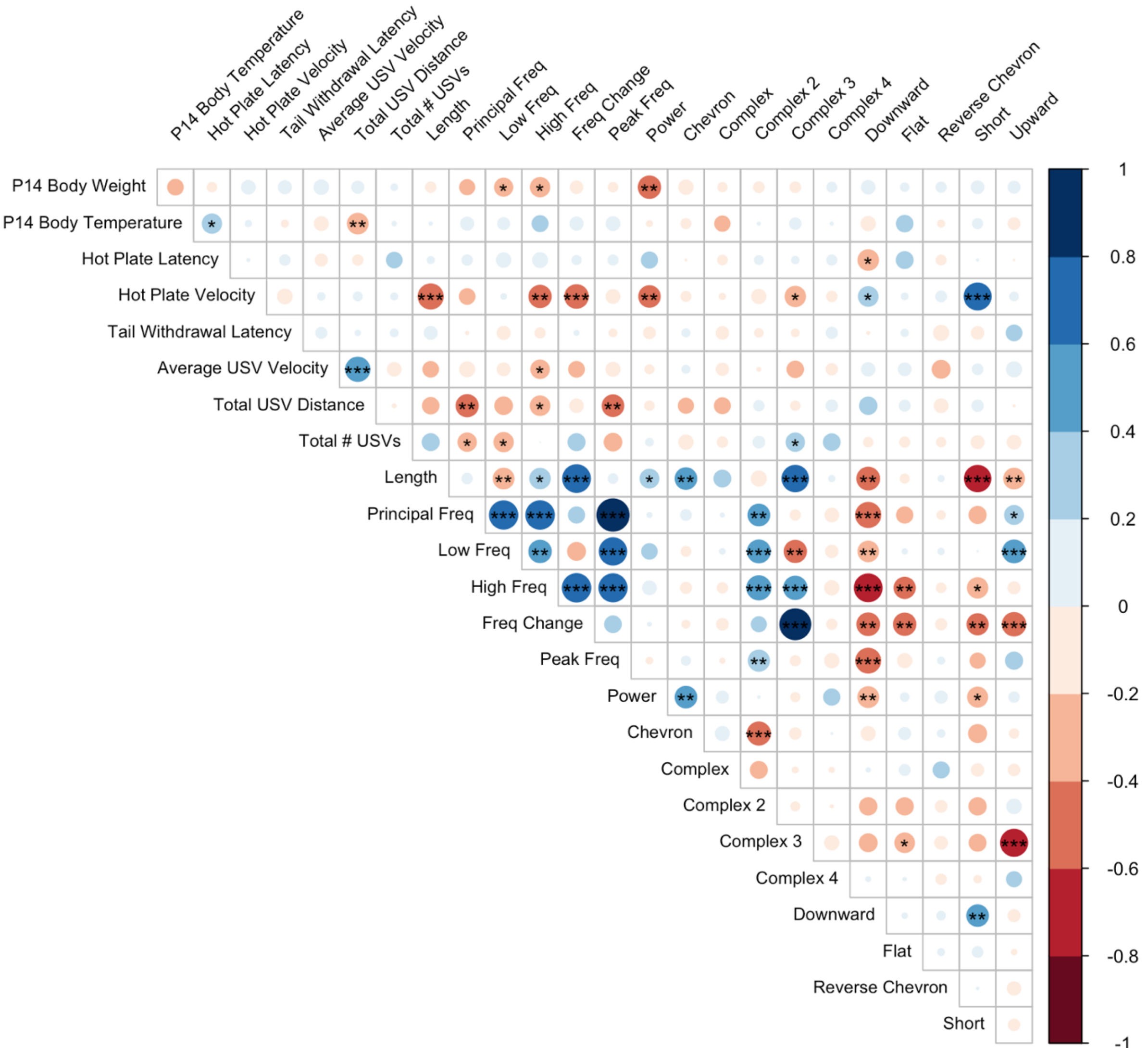
